# Combinatorial effect of miR-486-5p & miR-22-3p mimics thrombopoietin’s impact on hematopoietic stem & progenitor cells

**DOI:** 10.1101/2020.02.09.941104

**Authors:** Chen-Yuan Kao, Jinlin Jiang, Eleftherios T. Papoutsakis

## Abstract

Megakaryocytes shed and release submicron size microparticles (MkMPs), the most abundant microparticle in circulation. We have previously reported that MkMPs target peripheral-blood CD34^+^ hematopoietic stem/progenitor cells (HSPCs) to induce megakaryocytic differentiation and proliferation, and that small RNAs delivered to HSPCs via MkMPs play an important role in the development of this phenotype. Here, using single-molecule real-time (SMRT) RNA sequencing (RNAseq), we identify the top seven most abundant microRNAs (miRs) in MkMPs as potential candidates in mediating the effect of MkMPs on HSPCs. Using miR mimics, we demonstrate that among the seven most abundant miRs, two, miR-486-5p and miR-22-3p, are able to drive the Mk differentiation of HSPCs in the absence of thrombopoietin (TPO). The effect of these two miRs is comparable to the TPO- or MkMP-induced megakaryocytic differentiation of HSPCs, thus suggesting that these two miRs are responsible for this MkMP-induced phenotype. To probe the signaling through which MkMPs might enable this phenotype, we used kinase inhibitors of potential signaling pathways engaged in megakaryocytic differentiation. Our data suggest that MkMP-induced Mk differentiation of HSPCs is enabled through JNK and PI3K/Akt/mTOR signaling. Our data show that MkMPs activate Akt and mTOR phosphorylation. Furthermore, MkMPs downregulate PTEN expression, a direct target of miR-486-5p and a negative regulator of PI3K/Akt signaling, via JNK signaling. Taken together, our data provide a mechanistic understanding of the biological effect of MkMPs in inducing megakaryocytic differentiation of HSPCs, which, as was previously suggested, is a phenotype of potential physiological significance in stress megakaryopoiesis.

**Key Points:** - miR-486-5p and miR-22-3p drive megakaryocytic differentiation in the absence of thrombopoietin.
- Megakaryocytic microparticles trigger megakaryocytic differentiation of CD34^+^ cells through JNK and PI3K/Akt/mTOR signaling.

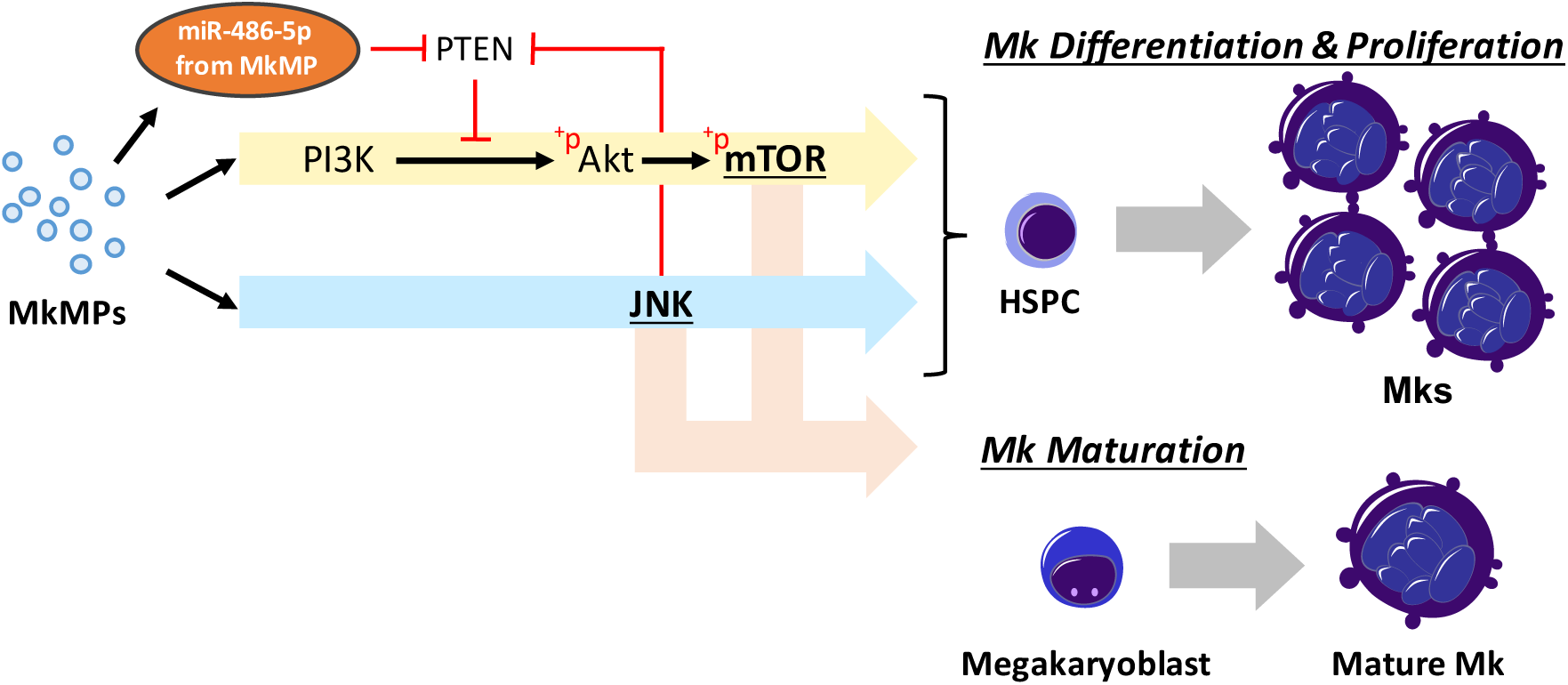
Visual Abstract

## Introduction

Cell-derived microparticles (MPs) are sub-micron size (0.1-1.0 µm) extracellular vesicles (EVs) that play an important role in cell-to-cell communication by carrying and transferring native cargo, including proteins, lipids, and RNAs to target cells.^1,2^ Cargo delivery triggers the development of complex phenotypes through mechanisms involving signaling and, broadly, regulation of gene expression.^3-5^

miRs are small non-coding RNAs regulating gene expression at the post-transcriptional level by targeting specific mRNAs leading to mRNA degradation or translational inhibition.^6^ EV-mediated transfer of miRs between cells has been studied in various cell types.^7,8^ Single-molecule real-time (SMRT) RNA sequencing (RNAseq) was used to identify specific RNAs involved in EV-triggered phenotypes of target cells.^9,10^ In these and other studies, a single EV miR was identified as responsible for the biological phenotype. There is increasing evidence however that two or more miRs are involved in co-regulating the same biological program or process in cancer^11,12^ and normophysiology.13 Combinations of miRs have been also used synthetically to regulate biological processes.^13,14^ However, cooperation between miRs from an EV has been rarely examined. Recently, Xu *et al* reported that a group of EV-associated miRs likely mediates pro-inflammatory cytokine production in a murine sepsis model,^15^ but mechanistic understanding was not pursued.

We have previous shown that megakaryocytic microparticles (MkMPs) can induce Mk differentiation of mobilized peripheral blood CD34^+^ Hematopoietic Stem and Progenitor cells (HSPCs; the term used henceforth to refer to these cells)^16,17^ in the absence of exogenous Thrombopoietin (TPO), yet similar in potency and effect as TPO. MkMPs are highly enriched in small RNAs. RNase treatment, differentially depleting the small RNA pool, attenuated the ability of MkMPs to trigger Mk differentiation of HSPCs,^16^ thus suggesting that small RNAs, possibly miRs, can mimic TPO signaling in HSPCs, a hitherto unknown possibility and mechanism. TPO-induced signaling starts with the binding of TPO to its receptor c-Mpl, which activates Janus-family kinases (Jaks).^18,19^ Downstream signaling pathways include signal transducers and activators of transcription (STAT),^20^ mitogen-activated protein kinases (MAPKs), and notably phosphoinositide 3-kinase (PI3K)/Akt/mammalian target of rapamycin (mTOR).^21^ Within the MAPK family, the MEK-ERK1/2 (MAPK kinase-extracellular signal-related kinases 1 and 2) signaling has been shown to play an important role in TPO-induced Mk development,^22^ while p38-MAPK was shown to be involved in TPO-mediated hematopoietic stem cell (HSC) expansion^23^ and erythropoiesis.^24^ Although TPO have been shown to activate c-Jun amino-terminal kinases (JNKs) signaling,^25^ there is no known role for JNK in TPO-induced Mk development.

Here, we tested the hypothesis that MkMP miRs are responsible for the observed phenotype and aimed to identify such possible miRs. Using SMRT RNAseq of the small-RNA content of MkMPs, platelet-like particles (PLPs), their parent Mk cells, and platelets (PLTs), we identified seven (7) miRs highly enriched in MkMPs, and show that among those, the combination of two miRs is capable of generating a phenotype similar to that of MkMPs or TPO in inducing Mk differentiation of HSPCs. In order to pursue possible signaling mechanisms by which MkMPs mimic TPO signaling, we used kinase signaling inhibitors in MkMP-induced Mk differentiation of CD34^+^ HSPCs and identified two key pathways (JNK and PI3K/Akt/mTOR) as important in this phenotype.

## Materials and Methods

### Generation of Megakaryocytic MPs (MkMPs) from cultured Megakaryocytes (Mks)

CD34^+^-derived Mks were cultured as described^26^ starting with frozen G-CSF-mobilized human peripheral blood CD34^+^ cells (Fred Hutchinson Cancer Research Center). MkMPs were isolated from the culture medium of the day-12 Mk culture as described.^16^

### Transfection of CD34^+^ HSPCs with miR mimics

200,000 CD34^+^ cells were freshly thawed and cultured in IMDM supplemented with 20% BIT 9500, and 100 ng/mL SCF. After 3 hours, cells were transfected with 8 µM of miR mimics, non-targeting miR (miR-NC), or without miR (No miR) using the Amaxa Nucleofector II with program U-08. After transfection, cells were cultured in IMDM supplemented with 10% BIT 9500, 50 ng/mL SCF, and 1 ng/mL IL-3, without TPO. Cells cultured in TPO-supplemented medium (100 ng/ml TPO), or co-cultured with MkMPs served as positive controls (TPO, MkMP). The medium was replaced one day after transfection. At days 7, 10 and 13, cells were harvested for flow-cytometric analysis of CD41, CD42b expression, and Mk (CD41+-cell) and total cell measurements. At day 13, cells were harvested for serotonin (5-HT), von willebrand factor (vWF), beta 1 tubulin (TUBB1), and DAPI staining, as described^17^. The images were taken by ZEISS LSM 880 multiphoton confocal microscope. At day 16, cells were harvested for ploidy analysis by flow-cytometric analysis as described^27^.

### miR-inhibitor experiments

Loading of miR inhibitors into MkMPs was performed as described^28^. Briefly, MkMPs were loaded with 8 µM of miR-486-5p inhibitor, or miR-22-3p inhibitor by electroporation. 600,000 CD34^+^ HSPC cells were freshly thawed, followed by the co-culture with MkMPs or miR-inhibitor-loaded MkMPs (30 MPs per cell), or vehicle control in IMDM supplemented with 10% BIT 9500 and 50 ng/mL SCF. Cells were harvested for flow cytometric analysis of CD41 expression at day 4, 7, and10. Total cell and Mk cell numbers were measured at day 10 of co-culture.

### Signaling-inhibitor experiments

60,000 CD34^+^ HSPCs were pretreated with signaling inhibitors for 30 min, followed by co-culture^16, 17^ with MkMPs (30 MPs per cell) in IMDM supplemented with 10% BIT 9500 and 50 ng/mL SCF. Inhibitors were replenished at days 3 and 7. At day 4, 7 and 12, cells were harvested for flow cytometric analysis of CD41, CD42b, and CD34 expression. Total-cell and Mk counts were measured at day 7. Inhibitor concentrations and treatment times were based on published studies.^29, 30^

Other experimental details can be found in Supplemental Materials & Methods. They include: Generation of Megakaryocytic MPs (MkMPs) from cultured Megakaryocytes (Mks) starting with CD34^+^ HSPCs, Chemicals and Reagents, Isolation of platelet-like particles (PLPs) Megakaryocytic MkMPs, Human platelets, RNA extraction and library preparation for RNAseq analysis, RNAseq data analysis, Immunoblotting, Quantitative reverse transcription PCR (qRT-PCR), Intracellular protein analysis by flow cytometry, and statistical analysis.

## Results

### The miR content of huMkMPs is well preserved among donors, with seven miRs making up 57% of the total miR content

We have previously shown that MkMPs were enriched in small RNAs,^16^ which play an important role in triggering Mk differentiation of HSPCs.^16, 17^ We hypothesized that miRs are the dominant MkMP components in inducing and promoting Mk differentiation. To characterize and carry out comparative analysis of the MkMP miR profile, total RNA was extracted from day-12 cultured Mks (starting from CD34^+^ cells of 3 donors), MkMPs and platelet-like particles (PLPs) generated from the corresponding day-12 cultured Mks. For comparison, we also extracted and analyzed RNA from human platelets (PLTs; 2 donors unrelated to the CD34^+^-cell donors). Small RNA libraries were prepared from extracted RNA for SMRT RNAseq analysis. At the average miR expression level (Count Per Million, CPM) of ≥ 1, RNAseq identified 514, 609, 589, or 484 miRs in Mks, MkMPs, PLPs, or PLTs, respectively. To identify highly expressed miRs, we used CPM ≥ 1000 as a criterion, and identified 63 miRs as highly abundant (accounting of 96.1% of total miR content). The Venn diagram (Figure 1A) shows that Mks and MkMPs share 491 miRs, while 446 miRs were shared between MkMPs with PLTs, or PLPs with PLTs. The miR MkMP profile was most distant from that of PLTs and Mks in decreasing distance order, and closest to the PLP miR profile. The miR profiles from 3 donors for Mks, MkMPs and PLPs, or two PLTs are consistent and reproducible (Figure S1). RNAseq analysis also identified other small RNAs, such as Piwi-interacting RNA (piRNA, 18-40 nt, Table S1), small nucleolar RNA (snoRNA, 40-150 nt, Table S2) or non-snoRNA (Table S3).

**Figure 1.**
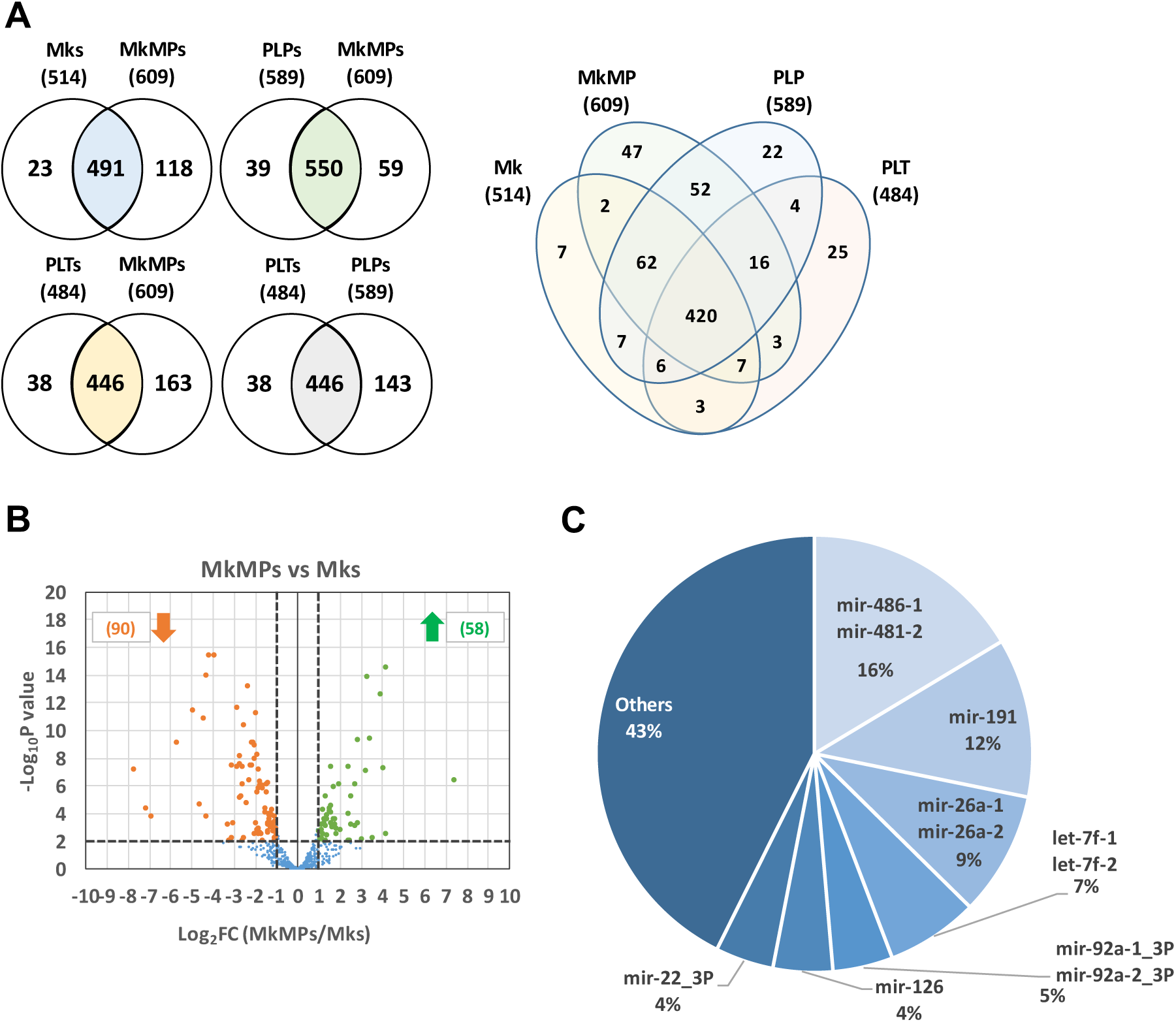
RNAseq analysis of miRs. (A) Venn diagram showing the number of miRs detected in (CPM ≥ 1) and shared among Mks, MkMPs, PLPs, and PLTs. (B) Volcano plot showing the results from the differential expression analysis of miRs in MkMPs vs. Mk cells. 58 miRs were significantly (*p < 0.01*) enriched (fold change ≥ 2) in MkMPs, while expression levels of 90 MkMP miRs were significantly lower (fold changes ≤ 0.5) than in Mk cells. (C) Pie chart showing the miR distribution in MkMPs. The top seven (7) miRs are shown in detail.

Next, we carried out differential expression analysis of MkMP miRs against the miRs in Mk cells, PLPs and PLTs. The hypothesis was that this analysis might identify miRs that could mediate the ability of MkMPs to induce Mk differentiation of HSPCs, assuming that such miRs were also abundant in MkMPs. Figure 1B summarizes the differential miR-expression analysis between MkMPs and Mk cells. Table S4 lists the 18 miRs which are highly enriched in MkMPs and their CPM≥100. Only 2 miRs, mir-19b-1//mir-19b-2_3p and mir-181b-1//mir-181b-2 were within the top 50 most abundant miRs in MkMPs. Since each of the highly-enriched miRs (Table S4) only accounts for less than 0.2% of total miR content in MkMPs, we reasoned that these miRs are not likely mediators of the observed phenotype.

Next, we hypothesized that highly abundant miRs in MkMPs would be more likely to achieve the observed biological phenotype. Table 1 lists the top 10 most abundant miRs in MkMPs, and for comparative purposes, also those of Mks, PLTs and PLPs. The top 20 most abundant MkMP miRs account for 81.8% of total miR count, while the top 7 miRs account for more than 57% of the total miR content (Figure 1C). Among the top 7 miRs in MkMP, only miR-22 is known to be involved in megakaryopoiesis by regulating the balance between erythroid and megakaryocytic differentiation *in vivo*.^31^ In miR-22 knock-out mice, megakaryopoiesis was enhanced after infection with lymphocytic choriomeningitis virus, while erythropoiesis was suppressed.^31^ Although there is no known role for miR-486-5p in megakaryopoiesis, miR-486-5p plays a role in CD34^+^-cell proliferation and erythroid differentiation^32^.

**Table 1.** Top 10 most abundant miRNAs in MkMPs, Mk cells, PLTs, or PLPs

| Mks |  |  |  | MkMPs |  |  |
| --- | --- | --- | --- | --- | --- | --- |
| Rank | GeneID | CPM | % | miRNA (Gene ID) | CPM | % |
| 1 | mir-191 | 231441 | 23.1 | mir-486-1//mir-486-2 | 164087 | 16.4 |
| 2 | mir-486-1//mir-486-2 | 195803 | 19.6 | mir-191 | 117782 | 11.8 |
| 3 | mir-99b | 164315 | 16.4 | mir-26a-1//mir-26a-2 | 91774 | 9.2 |
| 4 | mir-146b | 65671 | 6.6 | let-7f-1//let-7f-2 | 68251 | 6.8 |
| 5 | mir-26a-1//mir-26a-2 | 39625 | 4.0 | mir-92a-1//mir-92a-2_3P | 44480 | 4.4 |
| 6 | let-7f-1//let-7f-2 | 23623 | 2.4 | mir-126 | 44218 | 4.4 |
| 7 | mir-126 | 22642 | 2.3 | mir-22_3P | 43152 | 4.3 |
| 8 | mir-146a | 21993 | 2.2 | mir-21 | 38617 | 3.9 |
| 9 | mir-21 | 20197 | 2.0 | mir-146b | 34455 | 3.4 |
| 10 | mir-148a_3P | 17054 | 1.7 | mir-181a-2//mir-181a-1 | 33752 | 3.4 |
|  | Other miRNAs |  | 19.8 | Other miRNAs |  | 31.9 |
| PLTs |  |  |  | PLPs |  |  |
| Rank | GeneID | CPM | % | GeneID | CPM | % |
| 1 | mir-191 | 263871 | 26.4 | mir-191 | 162495 | 16.2 |
| 2 | mir-486-1//mir-486-2 | 80546 | 8.1 | mir-486-1//mir-486-2 | 146874 | 14.7 |
| 3 | let-7f-1//let-7f-2 | 76283 | 7.6 | mir-26a-1//mir-26a-2 | 84857 | 8.5 |
| 4 | mir-99b | 71050 | 7.1 | let-7f-1//let-7f-2 | 64288 | 6.4 |
| 5 | mir-10a | 54139 | 5.4 | mir-146b | 49986 | 5.0 |
| 6 | mir-26a-1//mir-26a-2 | 51712 | 5.2 | mir-126 | 43506 | 4.4 |
| 7 | mir-146a | 43647 | 4.4 | mir-99b | 42826 | 4.3 |
| 8 | mir-92a-1//mir-92a-2_3P | 43032 | 4.3 | mir-21 | 40419 | 4.0 |
| 9 | mir-146b | 25104 | 2.5 | mir-22_3P | 32727 | 3.3 |
| 10 | mir-181a-2//mir-181a-1 | 23932 | 2.4 | mir-181a-2//mir-181a-1 | 25520 | 2.6 |
|  | Other miRNAs |  | 26.7 | Other miRNAs |  | 30.7 |

### miR-486-5p in combination with miR-22-3p largely recapitulates the megakaryopoietic effect of TPO and MkMPs on CD34^+^ HSPCs

Based on previously unpublished miR data presented above, using miR mimics,^28^ we have recently demonstrated a role of miR-486-5p in Mk differentiation of CD34^+^ HSPCs.^28^ To determine if there is a dose effect of miR mimics on Mk differentiation of HSPCs, using CD41 expression as the key early Mk-differentiation marker, we carried out a pilot study with two donor CD34^+^-cell samples. CD34^+^ HSPCs were transfected with high (8 µM) or low (2 µM) concentrations of miR-486-5p or miR-22-3p mimics were and cultured without TPO. At day 10 of the culture, 33.8% or 27.0% of cells transfected with 8 µM miR-486-5p or 8 µM miR-22-3p were CD41^+^, while only 26.1% or 23.7% were CD41^+^ when transfected with 2 µM miR-486-5p or 2 µM miR-22-3p, respectively (Figure S2A), thus indicating a dose effect of miRs on HSPC Mk differentiation. An example of flow cytometric analysis histograms for selecting CD41^+^ population based on proper IgG control were shown in Figure S2B. Since CD41 and CD61 form a complex (gpIIb/IIIa), the expression level of CD41 and CD61 are virtually identical (Figure S2C). Based on published literature, miR concentrations from 1 nM to 3.6 µM have been used to transfect HSPCs.^33–35^ In the following studies, since the seven MkMP miRs we examined are highly abundant in MkMPs, and due to the fact that multiple MkMPs are taken up by a single recipient CD34^+^ HSPC,^16^ we hypothesized that higher concentrations of these seven miRs are likely delivered to CD34^+^ HSPC via MkMPs. Therefore, we chose 8 µM of miR mimics for the following experiments.

Focusing on the top 7 most abundant MkMP miRs (Figure 1C), to identify the most likely miR(s) that might impact Mk differentiation of CD34^+^ cells, we directly transfected 200,000 CD34^+^ HSPCs with 8 µM of each miR mimic separately. Transfected cells were cultured in IMDM supplemented with 10% BIT and 50 ng/ml SCF, but without TPO. Expression of CD41 and total Mk and total cell counts were examined at day 13 and 10. Negative controls were CD34^+^ cells exposed to same electroporation conditions without any miR, or with negative control miRs (miR-NC). Positive control was CD34^+^ cells exposed to same electroporation conditions cultured with 100 ng/ml TPO. Among the top 7 MkMP miRs, at day 13, compared to “miR-NC”, miR-486-5p significantly induced and promoted Mk differentiation of CD34^+^ HSPCs, achieving the highest percent (38.3%) of CD41^+^ cells, approaching that of the TPO control (Figure S3A). The miR-22-3p mimic significantly enhanced cell expansion by up to 72% or 61% (Figure S3B) compared to “No miR” or “miR-NC” controls, respectively. These results suggest that miR-486-5p plays a role in Mk differentiation, while miR-22-3p promotes total cell proliferation.

Combinations of small RNAs (siRNAs or miRs) have been shown to improve cell proliferation,^14^ and alter cellular phenotypes.^13^ We thus hypothesized that MkMP-induced Mk differentiation of CD34^+^ HSPCs might be mediated by miR-486-5p and miR-22-3p acting together. We thus examined their combinatorial targeting on CD41^+^ or CD42b^+^ expression, Mk-cell count, and total cell count in TPO-free cultures post transfection of CD34^+^ cells. Co-culture of CD34^+^ HSPCs with MkMPs, or CD34^+^ HSPC culture supplemented with TPO served as positive controls; all CD34^+^ cells were exposed to the electroporation conditions used for miR transfection, which, as would be expected, would lead to attenuated culture outcomes in terms of Mk differentiation and expansion. miR-486-5p significantly promoted Mk differentiation of CD34^+^ HSPCs with 41% of cells expressing CD41, while 43% of cells co-transfected with miR-486-5p and miR-22-3p were CD41^+^ (Figure 2A, 2B). Notably, compared to positive controls (MkMP, TPO), which resulted in 60% or 56% of the cells expressing CD41 by day 13, the miR-486-5p mimic alone achieved ca. 70% of their effect. Cells in each condition were also examined for expression of CD42b (a late Mk marker) at day 10. miR-486-5p or the combination of miR-486-5p and miR-22-3p mimics significantly enhanced CD42b expression with 20% and 22% of cells expressing CD42b (Figure 2C), indicating that miR-486-5p mediates Mk maturation. Representative quadrant plots (Figure 2D) demonstrate the combinatorial effect of miR-486-5p and miR-22-3p on CD41^+^CD42b^+^ expression (upper-right quadrant, 29.2% double positive) at day 13, which is ca. 2 and 3 times higher than the “No miR” and “miR-NC” negative controls, respectively, and closer to that of the two positive controls (MkMP, 35.2%; TPO, 38.9%)(the percent of CD41^+^CD42b^+^ cells at days 10 and day 13 are plotted in Figures S4A & S4B). Moreover, miR-486-5p targeting resulted in a significant increase (up to 98%) in the number of total Mks at day 13 compared to negative controls (Figure 2E). Compared to negative controls, the combinatorial effect of miR-486-5p and miR-22-3p resulted in a ca. 2.6-fold increase of total Mk cells at day 13, virtually matching the effect of TPO (Figure 2E).

**Figure 2.**
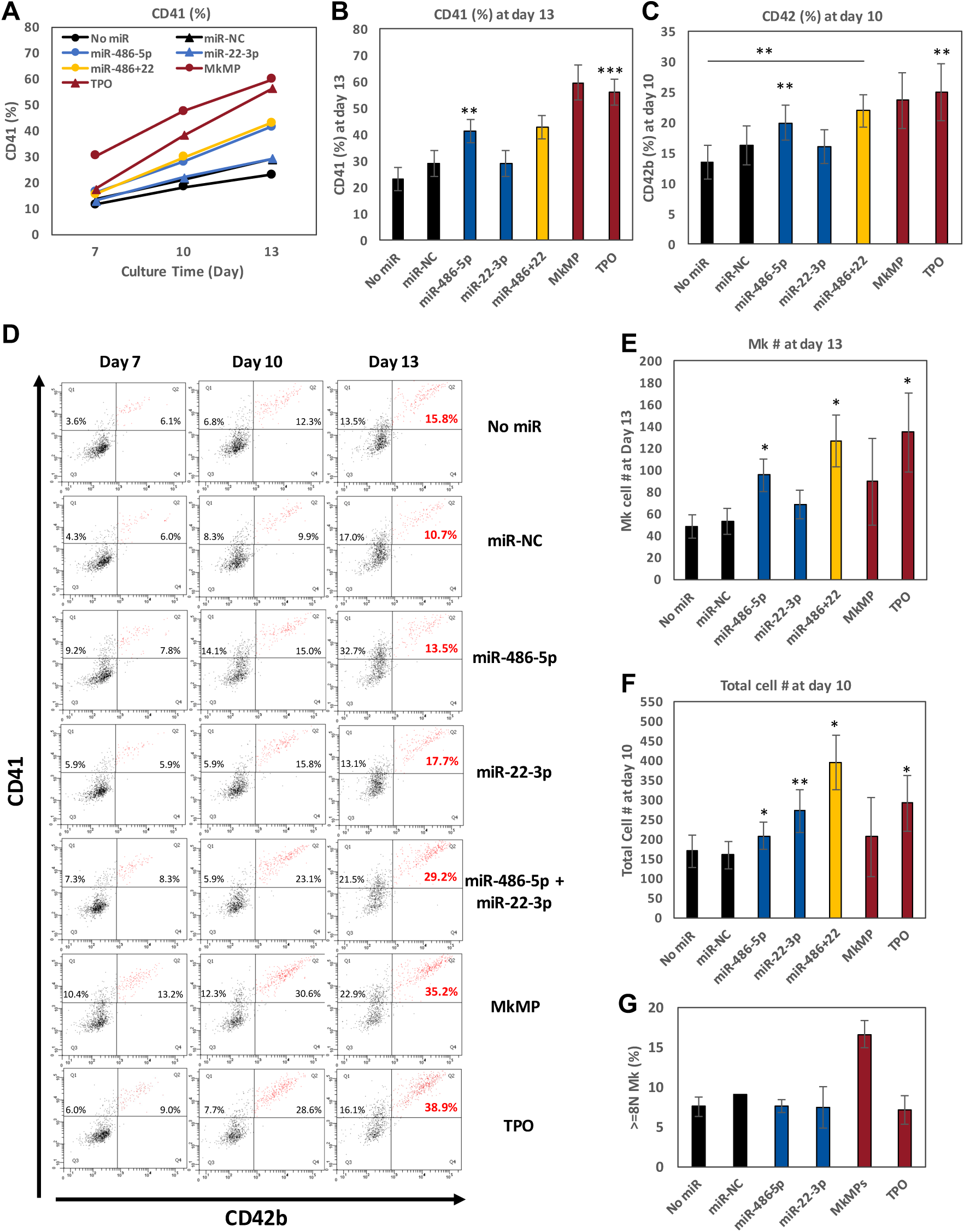
The effect of single or multiple miRs on Mk differentiation. CD34^+^ HSPCs were transfected with miR mimics (N=8), miR negative control (miR-NC, N=8), or without miRs (No miR, N=8), and cells were cultured in minimal medium (IMDM supplemented with 10% BIT and 50 ng/ml SCF) without TPO. Cells cultured in TPO-supplemented medium (100 ng/ml TPO, N=6) or cells co-cultured with MkMPs (N=3) served as positive controls (TPO, MkMP). Cells were harvested for flow cytometric analysis for (A, B) CD41 expression at days 7, 10 and 13, and (C) CD42b expression at day 10. (D) Representative flow cytometric dot plots for CD41 and CD42b expression at day 7, 10 and 13 with quadrant gates. (E) Mk (CD41^+^) cell numbers at day 13, or (F) total cell numbers at day 10 were measured by flow cytometry, respectively. (G) At day 16, cells were harvested for ploidy analysis by flow cytometric analysis. Error bars in (A, B, C, E, F, G) represent the standard error of mean from 3-8 biological replicates. Statistical comparison analysis was performed between each experimental group against the two negative controls (No miR or miR-NC) unless otherwise shown on panel C. *\*p < 0.05, **p < 0.01, ***p < 0.001*. Electroporation of CD34^+^ HSPCs damages and alters the properties of CD34^+^-cell membrane thus resulting in attenuated MkMP-induced Mk differentiation of CD34^+^ HSPCs (data not shown).

Similar to the pilot study, miR-22-3p significantly enhanced cell proliferation, with up to 71% of an increase on total cell numbers, compared to negative controls (Figure 2F, Figure S4C). The combinatorial targeting of miR-486-5p and miR-22-3p resulted in a 2.5-fold increase in total cell numbers, compared to negative controls (Figure 2F) and a 33% increase in total Mk cells compared to miR-486-5p alone (Figure 2E). These results demonstrate that combinatorial miRs targeting induces megakaryocytic differentiation in the absence of TPO, a novel and unexpected finding.

To further examine if miR-486-5p or miR-22-3p is capable of promoting late megakaryocytic differentiation of CD34^+^ HSPCs, we first examined polyploidization at day 16 for key experimental conditions (CD34^+^ transfected with miR-486-5p, miR-22-3p, or control No miR, miR-NC, as well as control CD34^+^ cells cultured post-electroporation with TPO or MkMPs) (Figure 2G). These data show that electroporation suppresses Mk polyploidization under all conditions, and that only MkMPs can partially rescue this suppression. Polyploidization is statistically identical for all other conditions. Next, we examined the cells at day 13 by confocal microscopy for expression of beta-1 tubulin (TUBB1), von Willebrand factor (vWF), and serotonin (5-HT), which are indicators of Mk maturation and platelet formation ^17^. The images look similar for the three conditions: miR-486-5p, miR-22-3p & TPO (Figure 3). Despite the suppressive effect of electroporation on polyploidization, both miR-486-5p-transfected and miR-22-3p-transfected cells displayed a few proplatelet-like structures (red arrows in Figure 3), vWF and 5-HT expression similar to that of the TPO-only culture. We also identified pre-demarcation membrane system (DMS) structures (white arrows in Figure 3) in miR-486-5p- and miR-22-3p-transfected cells, and cells from TPO culture, indicating megakaryocytic maturation. These results suggest that miR-486-5p and miR-22-3p impart megakaryocytic characteristics similar to those of TPO. Only 5-HT expression was slightly higher in the TPO-induced culture than “miR-486-5p” or “miR-22-3p” culture. The negative controls (No miR, miR-NC) displayed no 5-HT or VWF expression. Due to the suppressive effect of electroporation on polyploidization, the impact of miR-486-5p and miR-22-3p on HSPC development only into early megakaryopoiesis is inconclusive. It is possible that they may also impart megakaryocytic maturation. There is no literature that would suggest that CD41/CD42 expression does not lead to Mk maturation. Overall, these data suggest that miR-486-5p and miR-22-3p may be the key MkMP molecules through which MkMPs program CD34^+^ HSPC into Mk differentiation.

**Figure 3.**
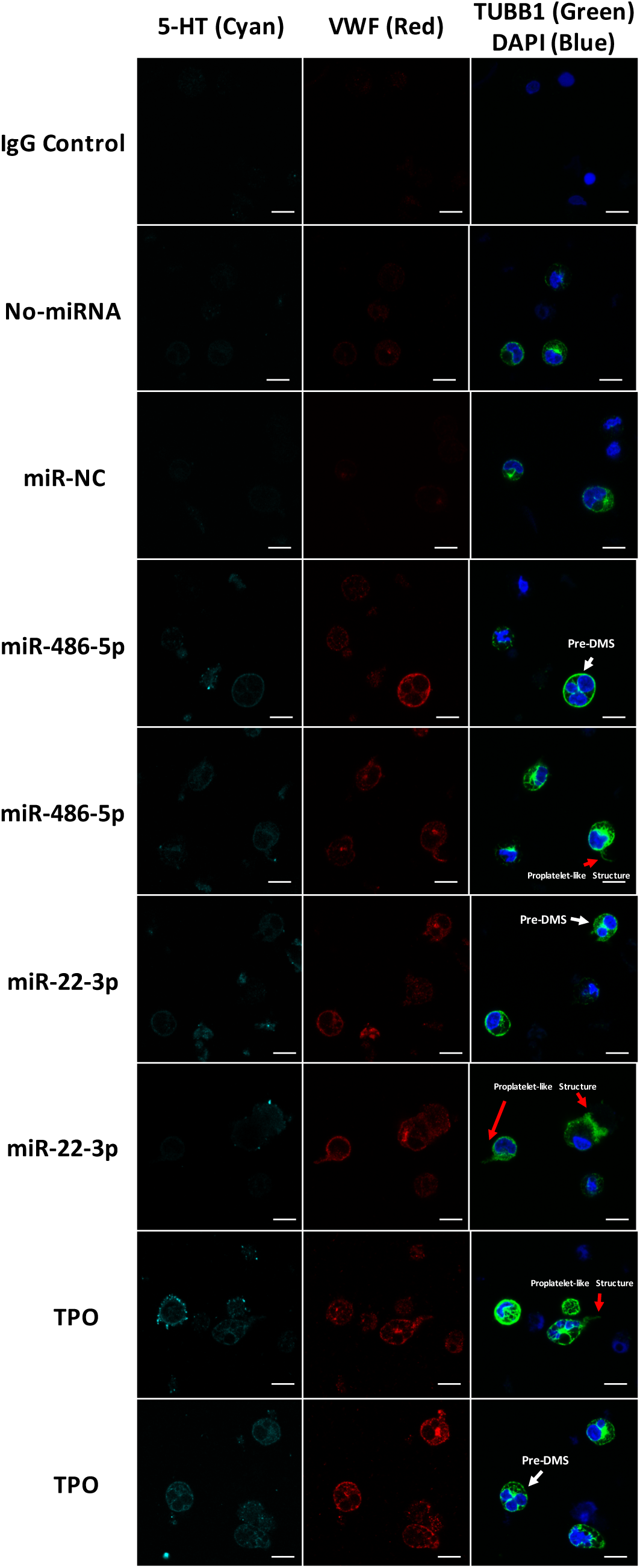
miR-486-5p and miR-22-3p enable Mk differentiation and maturation. CD34^+^ cells were transfected with miRNA mimics, non-targeting miRNA (miR-NC), or without miRNAs (No miR), and were cultured in minimal medium (IMDM supplemented with 10% BIT and 50 ng/ml SCF) without TPO. Cells cultured in TPO-supplemented medium (100 ng/ml TPO) served as positive control (TPO). Cells were harvested at day 13 and stained for 5-HT (cyan), vWF (red), TUBB1 (green), and DAPI (blue), and images were acquired by confocal microscopy. Scale bar: 10 µm. White arrows represent the pre-demarcation membrane system (DMS) structure. Red arrows represent the proplatelet-like structure.

### miR-486-5p in MkMPs serve as a key element mediating MkMP-induced megakaryocytic differentiation

Previously, we had shown (Figure 6 in Ref. ^28^) that co-culture of CD34^+^ cells with MkMPs loaded with exogenous miR-486-5p enhances megakaryocytic differentiation (22% higher fraction of CD41^+^ cells) compared to the co-culture with native MkMPs or MkMPs loaded with miR-NC. To further validate the importance of native miR-486-5p or miR-22-3p in MkMPs, here, we performed an experiment, where CD34^+^ cells were co-cultured with MkMPs loaded inhibitors of miR-486-5p or miR-22-3p (8 µM solutions for electroporation). Compared to MkMPs control, loading of miR-486-5p inhibitor to MkMPs significantly reduced the percentage of CD41^+^ cell by 14% (from 34.7% to 29.4%) at day 10, but the miR-22-3p inhibitor had no effect (Figure 4A). Since miR-486-5p is the most abundant miR in MkMPs (Figure 1), we hypothesized that higher levels of miR-486-5p inhibitors might be needed to achieve a stronger effect. Thus, when we doubled the concentration of miR-486-5p inhibitor for loading via electroporation MkMPs to 16 µM, it resulted in significantly lower number of total cells or Mk cells at day 10 (Figure 4B). Overall, these results further strengthen the evidence for a role of miR-486-5p in MkMPs in programming CD34^+^ HSPCs into Mk differentiation.

**Figure 4.**
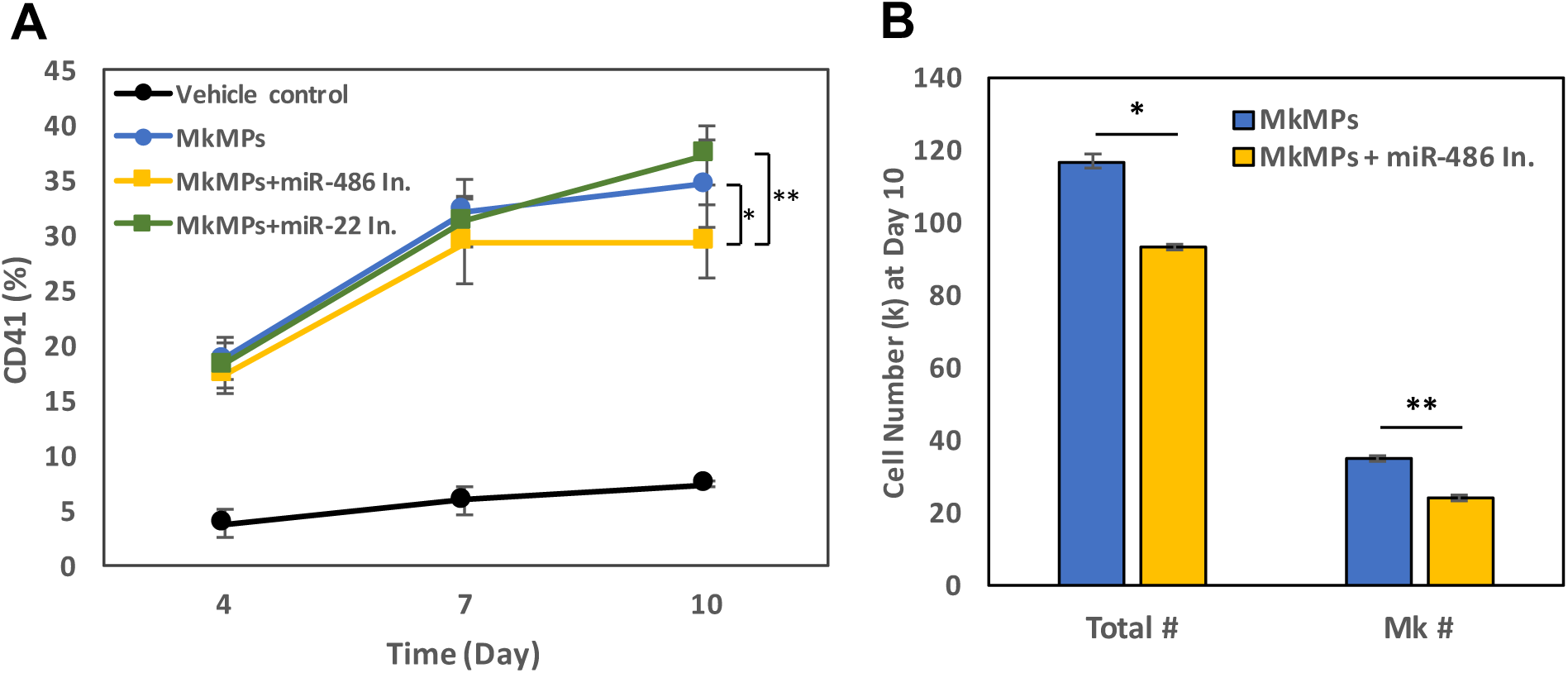
Loading of miR-486-5p inhibitor into MkMPs reduces the native effect of MkMPs in inducing Mk differentiation of CD34^+^ HSPCs. (A) CD34^+^ HSPCs were co-cultured with MkMPs, 8 µM-miR-486-5p inhibitor-loaded MkMPs, 8 µM-miR-22-3p inhibitor-loaded MkMPs, or vehicle control. Cells were harvested for flow cytometric analysis on CD41 expression at day 4, 7, and 10. (B) MkMPs, or 16 µM-miR-486-5p inhibitor-loaded MkMPs were co-cultured with CD34^+^ HSPCs. Total cell or Mk cell (CD41^+^ cells) numbers were measured at day 10.

### Use of signaling-pathway inhibitors suggests that JNK and PI3K/Akt/mTOR signaling regulate MkMP-induced Mk differentiation of HSPCs

To further investigate the effects of MkMPs in promoting megakaryocytic differentiation of CD34^+^ HSPCs, and how this might relate to the miR content of MkMPs, we next probed likely signaling pathways using kinase inhibitors. We started by examining signaling pathways known to be involved in TPO signaling as summarized on the Introduction. Briefly, CD34^+^ HSPCs were pretreated with kinase inhibitors of chosen signaling pathways (JNK, p38, MEK, PI3K, Akt and mTOR inhibitors, Table 2) for 30 min before they were co-cultured with MkMPs at the ratio of 10 MkMPs/cell. Inhibitors were replenished at days 3 and 7. CD41, CD42b and CD34 expression, and cell numbers were measured at days 4, 7, and 12. These kinase inhibitors are known to affect signaling by preventing activation (e.g., phosphorylation) of downstream molecules. Therefore, we expected that if a particular signaling pathway is involved in generating the phenotypic impact of MkMPs on CD34^+^ HSPCs, then we would observe reduced-phosphorylation of downstream molecules. Compared to MkMP control, the JNK (SP600125) and mTOR (rapamycin) inhibitors significantly suppressed Mk differentiation decreasing CD41 expression at day 7 from 44.3% (MkMP) to 32.8% (JNK) and 32.1% (mTOR), respectively (Figure 5A), while the p38 (SB203580), JNK, PI3K (LY-294002), or mTOR inhibitors significantly suppressed CD41 expression at day 12 (Figure 5B). CD42b expression was significantly inhibited by PI3K or mTOR inhibition (Figure 5C), suggesting that MkMPs promote Mk maturation via PI3K/mTOR signaling. JNK inhibition resulted in a higher fraction of cells expressing CD34 compared to MkMP or vehicle controls at both day 7 and day 12 (Figure 5D).

**Figure 5.**
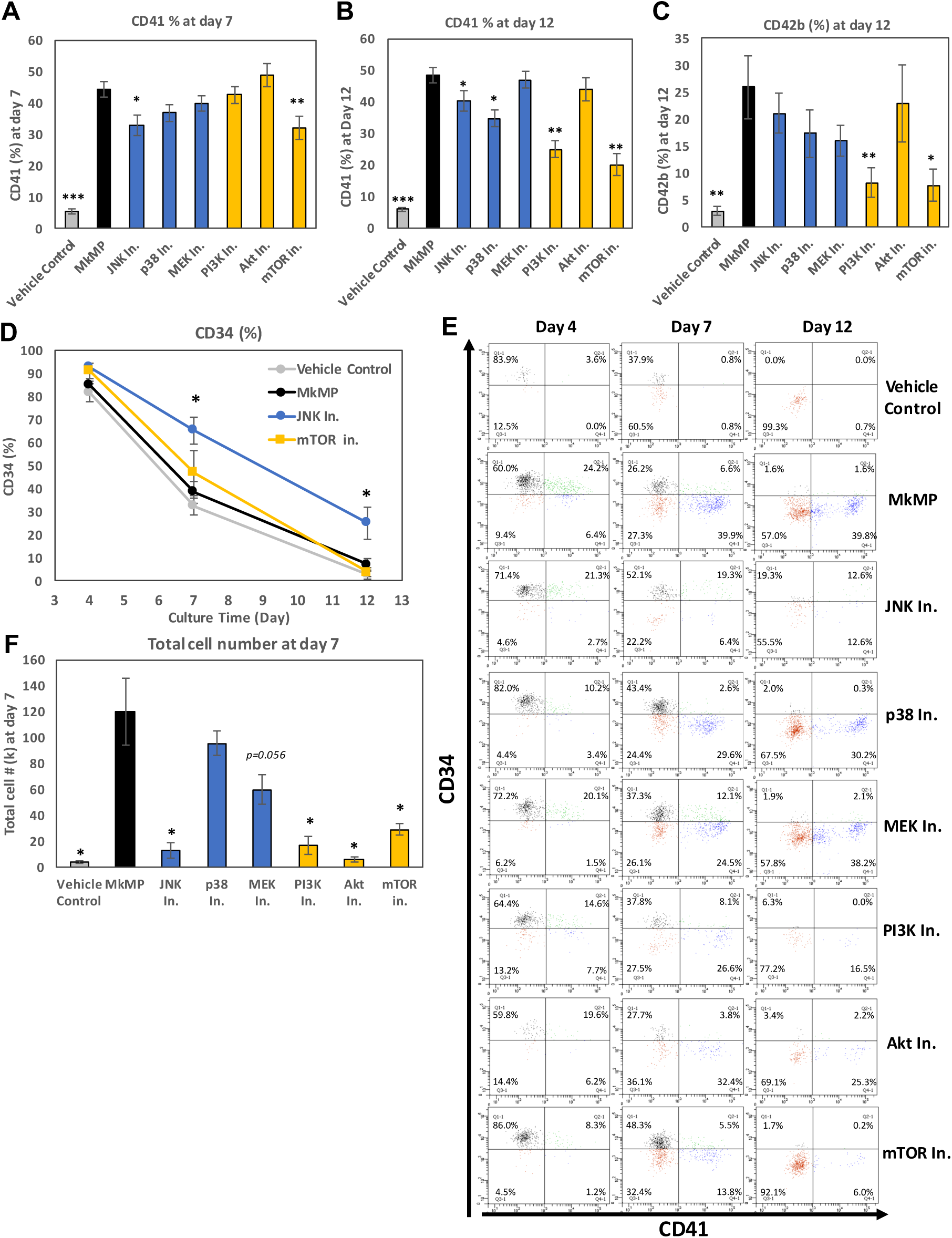
Impact of signaling inhibitors on MkMP-induced Mk differentiation. CD34^+^ HSPCs were pretreated with a signaling inhibitor, or solution without an inhibitor (MkMP control), and were co-cultured with MkMPs or vehicle control. Cells were harvested for flow cytometric analysis for (A, B) CD41 expression at day 7 and day 12, (C) CD42b expression at day 12, or (D) CD34 expression at days 4, 7, and 12. (E) Representative flow cytometric dot plots for CD34 and CD41 expression at day 4, 7 and 12 with quadrant gates. (F) Total cell numbers were measured at day 7. Error bars in (A-C) represent the standard error of mean of 6-8 biological replicates, while error bars in (D-F) represent the standard error of mean of 3 biological replicates \**p* < 0.05, \*\**p*<0.01 (compared to MkMP control)

**Table 2.** List of signaling inhibitors, their targets, and concentrations used

| Signaling Inhibitor | Target | Pathway | Concentration |
| --- | --- | --- | --- |
| SP600125 | JNK | MAPK | 10 uM |
| SB203580 | p38 |  | 10 uM |
| PD98059 | MEK |  | 10 uM |
| LY-294002 | PI3K | PI3K/Akt/mTOR | 10 uM |
| Wortmannin | Akt |  | 10 uM |
| Rapamycin | mTOR |  | 10 uM |

To further assess the impact of signaling inhibitors, representative flow-cytometric quadrant plots (Figure 5E) display the transition of MkMP-induced Mk differentiation of CD34^+^ cells, from CD34^+^CD41^-^ (upper-left) to CD34^+^CD41^+^ (upper-right), and on to CD34^-^CD41^+^ (lower-right). For example, by treating the MkMP-HSPC co-culture with JNK, p38, or mTOR inhibitors, at day 7, a larger fraction of cells was CD34^+^CD41^-^ (52.1%, 43.4%, and 48.3%, respectively) compared to MkMP control (26.2%), thus indicating that JNK, p38, or mTOR signaling is involved in the early MkMP-induced differentiation of CD34^+^ HSPCs. At day 12, 39.8% of cells were CD34^-^CD41^+^ in the MkMP control, while only 12.6%, 16.5%, 25.3%, or 6.0% of the cells were CD34^-^CD41^+^ using JNK, PI3K, Akt, or mTOR inhibitors. Treatment with the JNK, PI3K, Akt, or mTOR inhibitors significantly inhibited cell growth and decreased total-cell numbers by more than 75% at day 7 of co-culture (Figure 5F). Taken together, these data suggest that JNK, p38, PI3K, Akt and mTOR signaling are involved in MkMP-induced Mk differentiation of CD34^+^ HSPCs. PI3K and mTOR signaling appears to be involved in MkMP-promoted Mk maturation (Figure 5C), while JNK, PI3K, Akt, or mTOR signaling play an important role in MkMP-mediated cell proliferation (Figure 5G).

### MkMPs target JNK-mediated PI3K/Akt/mTOR signaling in HSPCs

Based on our findings above, both JNK and PI3K/Akt/mTOR appear to be involved in MkMP-induced cell proliferation and Mk differentiation (Figure 5). While PI3K/Akt signaling has been shown to be involved in TPO-mediated Mk differentiation, very little has been reported regarding JNK signaling in megakaryopoiesis.^18, 19, 21^ To pursue these findings further, we first examined expression of total and phosphorylated Akt and mTOR by immunoblotting. We expected lower phosphorylation levels of Akt or mTOR when using the Akt (Wortmannin) or mTOR inhibitor (Rapamycin), respectively. CD34^+^ HSPCs were pre-incubated with or without JNK, Akt, or mTOR inhibitors (Table 2), and co-cultured with MkMPs for 24 hours. The data (Figure 4A) show that total Akt expression and phosphorylated mTOR (p-mTOR) were higher (p-mTOR by 4.7 fold (Figure 6B)) in HSPCs co-cultured with MkMPs, but there were no changes on total mTOR levels. With the limited number of cells in CD34^+^ HSPCs cultures using signaling inhibitors, quantitating the very low expression levels of phosphorylated Akt became a significant challenge (data no shown). We thus examined total and phosphorylated Akt and mTOR levels using flow cytometry, which requires relatively few cells (Figure 6C-E). p-mTOR and Akt expression were significantly higher in the MkMP co-cultures, while treatment with JNK, Akt or mTOR inhibitors brought the expression levels back to that of vehicle control (Figure 6C & 6D). Total mTOR remained unaffected under all conditions (data not shown). Higher increase of p-mTOR levels was detected from immunoblotting (Figure 6B) than by flow-cytometric analysis (Figure 6C) likely because immunoblotting examines protein levels in both live and dead cells, while only live cells are examined by flow cytometry. Flow-cytometric analysis (Figure 6E) suggests that MkMPs also activate Akt phosphorylation. However, the level of p-Akt was not affected by the JNK inhibitor. These results suggest that, in CD34^+^ HSPCs stimulated by MkMPs, total Akt expression, but not total mTOR expression, is impacted by JNK signaling.

**Figure 6.**
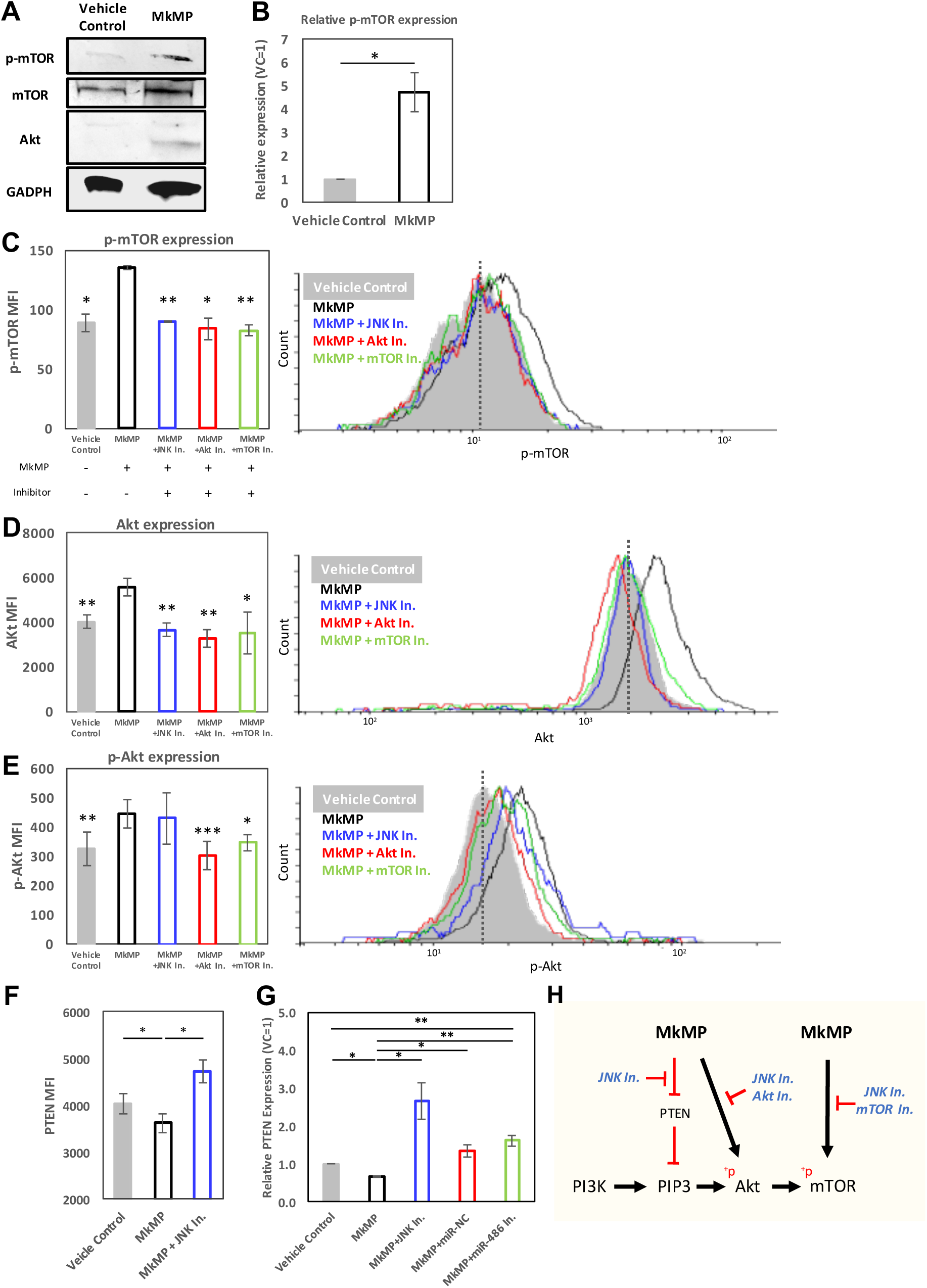
MkMPs activate PI3K/Akt/mTOR signaling in CD34^+^ HSPCs by targeting PTEN and mediated by JNK signaling. (A, B) CD34^+^ HSPCs were co-cultured with MkMPs or vehicle control. Cells were harvested after 24 hours of co-culture, and the level of targets including mTOR, p-mTOR, and Akt were examined by immunoblotting. (A) Representative immunoblot images out of 4 replicates. (B) Semi-quantification of p-mTOR expression from immunoblot images (n=4). (C-E) CD34^+^ HSPCs were pretreated with a signaling inhibitor (JNK, Akt, or mTOR inhibitor), or inhibitor vehicle (without an inhibitor), and were co-cultured with MkMPs or without (vehicle control). Cells were harvested after 24 hours of co-culture, and the level of p-mTOR, Akt, and p-Akt were examined and quantified by flow cytometric analysis (n=3-5). Representative histograms of each target are shown on the right. (F, G) CD34^+^ HSPCs pretreated with JNK inhibitor or solution without an inhibitor were co-cultured with MkMPs or vehicle control for 24 hours. (F) PTEN protein expression was quantified by flow cytometric analysis, while (G) *PTEN* mRNA levels were quantified by qPCR analysis. (H) Schematic diagram of signaling pathway triggered by MkMPs in CD34^+^ HSPCs, based on the results from (A-F). Error bars in (B) represent the standard error of mean of 2-4 biological replicates, while error bars in (C-G) represent the standard error of mean of 3-5 biological replicates. Statistical comparison analysis was performed between each experimental group against MkMP control (without inhibitor). \**p* < 0.05, \*\**p* < 0.01, \*\*\**p* < 0.001.

Phosphatase and TENsin homolog (PTEN) is a negative regulator of PI3K/Akt signaling in HSC development.^36, 37^ Loss of PTEN results in enhanced cellular proliferation^38^ and megakaryopoiesis^39^ due to overactive of PI3K/Akt signaling. We hypothesized that activation of PI3K/Akt/mTOR signaling by MkMPs might be PTEN mediated. We thus examined the mRNA level of *PTEN* by quantitative PCR, and PTEN protein level by flow cytometry and immunoblotting. The impact on PTEN mRNA levels appears higher than that on protein levels. Compared to the vehicle control, PTEN protein or mRNA levels were around 10% or 30% lower, respectively, in HSPCs co-cultured with MkMPs (Figure 6F, 6G). This effect was strengthened by the finding that the JNK inhibitor restored PTEN protein or mRNA levels by 17% or 260% above vehicle-control levels (Figure 6F, 6G, S5), respectively. These statistically-significant but relatively low-impact effects on PTEN protein expression would be expected given that the measurements were done within the first 24 hours of the co-culture, rather than late in the co-culture. Importantly, co-culture with MkMPs loaded with miR-486-5p inhibitor resulted in 2.4 or 1.2 fold higher PTEN-mRNA levels, compared to native MkMPs or MkMPs loaded with miR-NC, respectively (Figure 6G), thus suggesting that there is a relationship between miR-486-5p in MkMPs and PTEN in HSPCs.

To sum, our data suggest that MkMPs regulate Akt/mTOR signaling by enhancing Akt expression and activating Akt/mTOR, possibly mediated via JNK signaling including by PTEN targeting (Figure 6H).

## Discussion

### Synergistic action of two miRs mimicking TPO signaling

EV miRs are important mediators of EV-based cell-to-cell communication.^40^ Such miRs are either highly abundant or/and highly enriched in EVs.^9^ Here, we used RNAseq to identify miRs highly enriched in MkMPs, and examined the role of the most abundant miRs in promoting HSPC differentiation and cell proliferation (Figure 1C). As discussed, only few studies have examined combinatorial effects of two or more miRs on cell fate. Our results suggest that miR-486-5p induces Mk commitment and early development (CD41^+^ cells) of HSPCs (Figures 2A & 2E), miR-22-3p promotes cell proliferation (Figure 2F & Figure S4C) and Mk maturation (Figure 2D), and, combinatorically, Mk maturation (Figures. 2C & 2E), and Mk and total-cell expansion (Figures 2E & 2F). CD34^+^ transfected with miR-486-5p or miR-22-3p displayed Mk characteristics, including vWF/5-HT expression, DMS and proplatelet structures (Figure 3), similar to TPO-induced Mks. Together, miR-22-3p and miR-486-5p appear to mimic the TPO-induced Mk differentiation of CD34^+^ HSPCs with the notable outcome of Mk numbers being comparable to those in TPO-induced megakaryopoiesis (Figure 2E). This is the first report of two miRs inducing and co-regulating megakaryopoiesis of CD34^+^ HSPCs. It is possible that other abundant miRs (Figure S3), less abundant miRs, or other small RNAs (piRNA, snoRNA, non-snoRNA, Table S1-S3) may synergize with these two miRs to further promote Mk differentiation.

Several miRs have been previously reported as regulators of megakaryopoiesis.^33, 41–44^ These and all *in vitro* studies of miRs in megakaryopoiesis were carried out in the presence of TPO. To our knowledge, there have been no reports of miRs promoting megakaryocytic differentiation of CD34^+^ HSPCs in the absence of TPO.

miR-486-5p has recently been shown to regulate erythroid differentiation and survival of cord blood CD34^+^ cells via Akt signaling, both *in vitro* and *in vivo*.^32^ Conflicting roles for miR-22 have been reported in the development of hematopoietic malignancies, as a tumor suppressor^45^ or oncogenic.^46, 47^ Most recently, Weiss and Ito reported that miR-22 is upregulated during *in vivo* murine megakaryopoiesis, and that miR-22 knockout impairs megakaryocytic differentiation, while miR-22 overexpression promotes megakaryocytic differentiation in the K562 cell line.^48^ Their results suggest a similar miR-22 role as we report here (Figure 2D). miR-22-3p was also most recently shown to regulate mTOR signaling by targeting eukaryotic translation initiation factor 4E-binding proteins (eIF4EBP3) in human cervical squamous carcinoma cells.^49^

While miRs have been previously identified in human platelets, it is difficult to compare our RNAseq data (Table 1) to other studies, due to the fact that the data are collected and/or analyzed differently. For example, from microarray screening, miR-126, miR-197, miR-223, miR-24, and miR-21 were found to be the most highly expressed miRs in platelets^50^, which is different from our ranking of top miRs in platelets (Table 1). Nagalla *et al.* have also published a miR profile of human platelets from microarray analysis^51^. Depending on the tool used for miR analysis, the ranking of miRs in platelets varies significantly (Figure 3 in ^52^). The ranking of miRs in platelets from our data largely correlates with the data from the study of Kaudewitz & Skroblin (Figure 1 in ^53^), which used a similar strategy based on RNAseq. Lastly, Juzenas *et al.* provide a comprehensive miR dataset for leukocytes and erythrocytes.^54^

### JNK and Akt/mTOR signaling in MkMP-induced Mk differentiation of CD34^+^ HSPCs & a proposed model linking miR targeting to megakaryopoietic signaling

Multiple signaling pathways are engaged in TPO-induced megakaryopoiesis, including those of PI3K/Akt, MAPK, and Jak/STAT.^19, 55^ From the kinase inhibitor studies (Figures 5 & 6), we identified the role of JNK and PI3K/Akt/mTOR signaling. Although it has been shown that JNK can be activated by TPO,^25^ there is no known role of JNK signaling in Mk development.^55^ Using a JNK inhibitor in the co-culture of MkMPs with CD34^+^ HSPCs, CD34 expression was significantly maintained (Figure 5D) and Mk numbers at day 7 were significantly reduced, thus indicating that JNK signaling mediates early Mk differentiation and expansion in this MkMP-induced phenotype. Maintenance of CD34 expression is consistent with the recent finding that treatment of human cord blood CD34^+^ cells with JNK inhibitors (JNK-IN-8 or SP600125) enhanced the self-renewal of HSCs.^56^ We have also demonstrated that JNK signaling is involved in shear-induced Mk maturation and platelet production.^57^

mTOR is a major regulator of Mk development and maturation.^58^ Here, we showed that CD41 expression was significantly lower at days 7 and 12 in cells treated with an mTOR inhibitor prior to co-culture with MkMPs (Figure 5A & 5B). Akt expression was upregulated, and Akt and mTOR were phosphorylated in HSPCs co-cultured with MkMPs (Figure 6C-E), possibly through PTEN targeting. These results suggest that MkMPs activate Akt/mTOR signaling in HSPCs to induce Mk differentiation. Surprisingly, downregulation of PTEN expression by MkMPs was abolished upon treatment with a JNK inhibitor (Figures 6F, 6G & S5), together with reduced total Akt expression (Figure 6D) and mTOR phosphorylation (Figure 6C). These findings suggest that there is crosstalk between JNK and PI3K/Akt/mTOR signaling. A similar case has been previously reported, namely the negative regulation of PTEN by c-Jun, a downstream molecule in JNK signaling.^59^ To sum, our data suggest that it is possible that MkMP-induced Mk differentiation of CD34^+^ HSPCs is regulated by the circuit of JNK and Akt/mTOR signaling.

miR-486-5p have been shown to target PTEN and PI3K/Akt signaling in several cell types.^10, 60, 61^ Specifically, miR-486-5p regulates Akt signaling, cell proliferation and survival in cord-blood derived CD34^+^ cells by directly targeting PTEN.^32^ It is possible that miR-486-5p from MkMPs directly targets PTEN and activates PI3K/Akt/mTOR signaling in CD34^+^ HSPCs. Our data suggest that miR-22-3p plays an important role in cell proliferation and Mk maturation (Figure 2). The former is consistent with the finding that conditional miR-22 expression in the murine hematopoietic compartment increases hematopoietic stem cell self-renewal by directly targeting the tumor suppressor TET2.^47^ The latter is consistent with the finding that miR-22 promotes megakaryopoiesis by repressing the repressive transcription factor GFI1.^48^ The proposed model of Figure 7 captures and integrates our data and the current knowledge.

**Figure 7.**
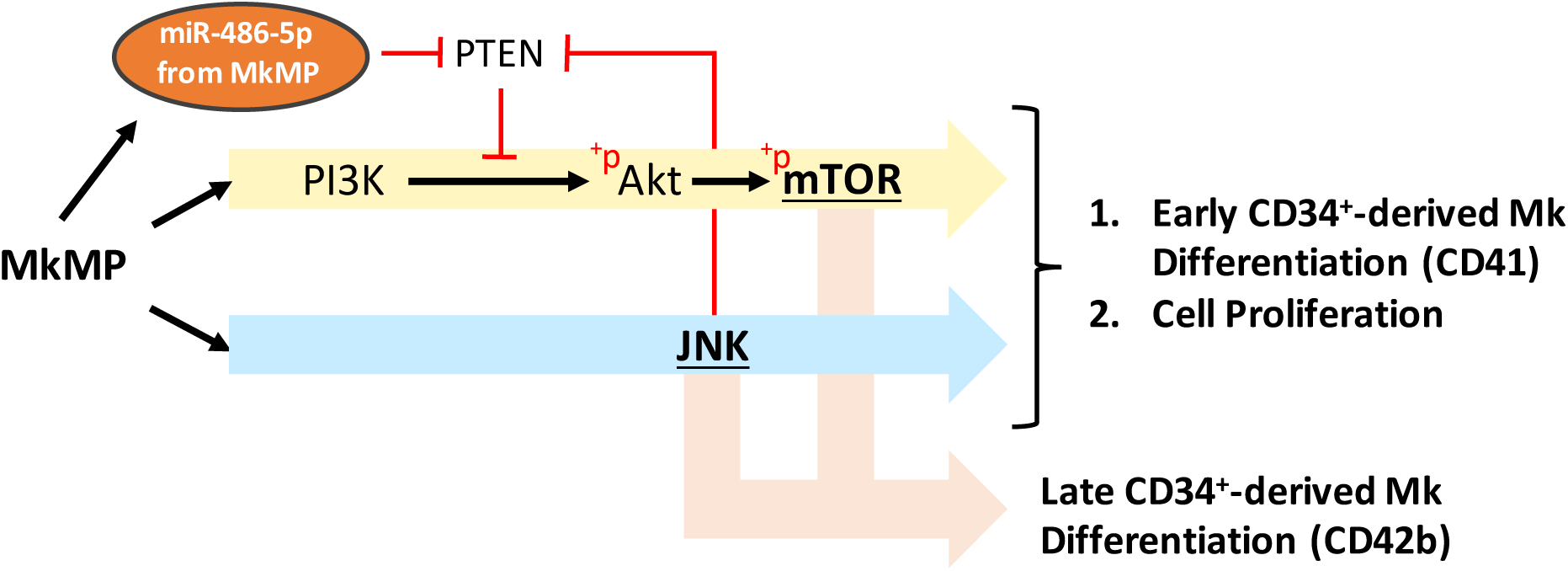
Schematic diagram of proposed model for MkMP-induced signaling in CD34^+^ HSPCs and its relationship to miRs in MkMPs. The model is based on the following pieces of evidence. Our data showed that co-culture of CD34^+^ HSPCs with MkMPs activates PI3K/Akt/mTOR and JNK signaling pathway toward cell expansion and megakaryocytic differentiation. Specifically, JNK and mTOR signaling are involved in late Mk differentiation (CD42b expression). A crosstalk between JNK and PI3K/Akt/mTOR was identified via the negative regulator of PI3K/Akt/mTOR signaling, PTEN. PI3K and PTEN were known major targets of miR-486-5p.

## Acknowledgements

We thank Bruce Kingham and members of DNA Sequencing and Genotyping Center (Univ. of Delaware) for assistance with RNAseq, and Shawn Polson and members of Bioinformatics Center (Univ. of Delaware) for assistance with differential analysis.

This project was supported by a grant (CBET-1804741) by the US National Science Foundation.

## Authorship Contributions

E.T.P. and CY.K and J.J designed the study and analyzed the data; CY.K. and J.J carried out the experiments. E.T.P. and CY.K. wrote the manuscript.

