## Supplemental material for "Combinatorial effect of miR-486-5p & miR-22-3p mimics thrombopoietin’s impact on hematopoietic stem & progenitor cells"

**Supplementary Data included:**

1. Supplemental Materials and Methods
2. Supplemental Tables: 4
3. Supplemental Figures: 5
4. Supplemental References: 8

**Supplemental Materials and Methods**

*Generation of Megakaryocytic MPs (MkMPs) from cultured Megakaryocytes (Mks) starting with CD34^+^ HSPCs*

CD34^+^-derived Mks were cultured as described*^1^* starting with frozen G-CSF-mobilized human peripheral blood CD34^+^ cells (Fred Hutchinson Cancer Research Center). Briefly, Cells were thawed and cultured in Iscove modified Dulbecco medium (IMDM, Gibco) supplemented with 20% BIT 9500 (Stemcell Tech.), 100 ng/mL TPO, 100 ng/mL stem cell factor (SCF), 2.5 ng/mL interleukin-3 (IL-3), 10 ng/mL IL-6 & IL-11 and human LDL under 5% O_2_ for 5 days. IL-3 was increased to 10 ng/mL and IL-6 was substituted with IL-9 at day 5. Cells were cultured under 20% O_2_ from day 5 to 7. At day 7, in order to achieve pure megakaryocyte culture, CD61^+^ cells were enriched by using MACS separation with anti-CD61 magnetic microbeads (Miltenyi). Enriched cells were then cultured in IMDM supplemented with 20% BIT 9500, 100 ng/mL TPO, 100 ng/mL SCF, and human LDL under 20% O_2_ for another 5 days. MkMPs were isolated from the culture medium of the day 12 Mk culture as described*^2^*.

*Chemicals and Reagents*

Recombinant human interleukin 3 (IL-3), IL-6, IL-9, IL-11, stem cell factor (SCF), and thrombopoietin (TPO) were purchased from PeproTech Inc. BIT 9500 was purchased from Stemcell Tech. Anti-CD61 magnetic microbeads and MACS cell-separation tools were purchased from Miltenyi. Fluorescein isothiocyanate (FITC)-conjugated anti-CD41, Phycoerythrin (PE)-conjugated anti-CD42b, allophycocyanin (APC)-conjugated anti-CD34, and IgG antibodies were purchased from BD bioscience. Signaling inhibitors, miRNA mimics, and miR-negative control were purchased from Sigma-Aldrich.

***Isolation of platelet-like particles (PLPs) Megakaryocytic Microparticles (MkMPs)***

PLPs and MkMPs were isolated as described *^2, 3^*. Briefly, cells and cell debris from day-12 CD34^+^-derived megakaryocyte culture, described above, were removed by centrifugation at 150 × g for 10 minutes. PLPs were collected from supernatant by centrifugation at 1000 × g for 10 minutes. MPs were then enriched via ultracentrifugation (Optima Max Ultracentrifuge and Rotor TLA55, Beckman Coulter) under 25,000 rpm for 30 min at 4 °C. After that, MPs were resuspended in IMDM or stored at -80 °C until used.

*Human platelets*

Blood for isolation of human platelets (PLTs) was collected*^3^* by venipuncture from adult healthy human volunteers after providing written informed consent as approved by the Institutional Review Board at the University of Delaware (IRB protocol # 622751). Briefly, 50 mL of blood was collected into a syringe with ACD buffer (trisodium citrate, 65 mM; citric acid, 70 mM; dextrose, 100 mM; pH 4.4) at a volume ratio of 1:6 (ACD:blood). Following that, blood was centrifuged at 250 × g for 10 min and the platelet-rich plasma was isolated from the supernatant. PLTs were then pelleted at 750 × g for 10 min, followed by 1 wash with HEN buffer (10 mM HEPES, pH 6.5, 1mM EDTA, 150 mM NaCl) containing 0.05 U/ml apyrase. After that, PLTs were resuspended in HEPES-Tyrode’s buffer (137 mM NaCl, 20 mM HEPES, 5.6 mM glucose, 1 g/l BSA, 1 mM MgCl2, 2.7 mM KCl, 3.3 mM NaH2PO4).

*RNA extraction and library preparation for RNAseq analysis*

11 small RNA libraries were prepared as described.*^4^* They include 3 biological samples of Mks, MkMPs, and PLPs, and 2 biological samples of human PLTs. Total RNA was isolated using the miRNeasy micro kit (Qiagen). RNA concentration was measured by NanoDrop (Thermo Scientific, ND1000) and size distribution of total RNA was analyzed using an ABI Prism 3130XL Genetic Analyzer at the University of Delaware (UD) Sequencing & Genotyping Center at the Delaware Biotechnology Institute. Small-size RNA (18-40 nt and 40-150 nt in size) was purified by 15% polyacrylaminde/urea gels and eluted from gels for library construction using Illumina TruSeq Small RNA Sample Prep kit according to the manufacturer’s protocol. Briefly, RNA was sequentially ligated with 3’ and 5’ adaptors, reverse transcribed to cDNA using SuperScript III reverse transcriptase (Invitrogen) and cDNA libraries were amplified by PCR Following that, a 6% polyacrylamide gel was used to purify cDNAs with size ranges of 140-160 base pairs (bp) and 160-275 bp derived from 18-40 nt and 40-150 nt input RNA, respectively. The 11 libraries described above were pooled together. 20 μL of pooled libraries at a final concentration of 10 nM was sequenced at UD’s Sequencing & Genotyping Center at Delaware Biotechnology Institute using 51 cycles on the Illumina HiSeq2500 DNA sequence analyzer.

*RNAseq data analysis*

Sequencing data analysis was provided by Dr. Shawn Polson and Jaysheel Bhavsar (Center for Bioinformatics & Computational Biology, UD). For small RNA (18-40 nt) sequencing data, a custom bioinformatics pipeline was used to end-trim raw reads to achieve an average quality score (Q) larger than 30, and to partition the data into miR and piRNA size fractions. Similarly, small RNA (40-150 nt) sequencing data were processed and mapped to small nucleolar RNA (snoRNA) and non-snoRNA. Only reads for which the flanking-adapter sequence was detected at the 3’ end were retained for analysis as they represent full-length sequencing of the molecule. Trimmed and filtered reads were then clustered if containing identical sequence and each cluster was aligned against human miR sequences downloaded from the miRBase (Release 21) *^5^*. miR reads were normalized by the number of counted reads per 1,000,000 total reads (Count per million, CPM). Differential expression analysis was performed using the edgeR Bioconductor Package *^6^*. The p-value was corrected by False Discovery Rate (FDR). Corrected p value <0.01 was used to define differentially expressed miR in MkMPs.

*Immunoblotting*

200,000 CD34^+^ cells were first pretreated with JNK inhibitor (or DMSO as a control) for 30 min, and co-cultured with MkMPs at 30 MPs/cell, or vehicle control for 16 hours. Immunoblotting was performed described *^7^*. Briefly, cells were lysed in NP-40 lysis buffer and proteins were separated by SDS-polyacrylamide gel electrophoresis using ExpressPlus 4-20% Bis-Tris polyacrylamide gels (Genscript #M42012) and the Mini-PROTEAN Tetra Vertical Electrophoresis Cell (Bio-Rad #1658004). Proteins were then transferred onto a nitrocellulose membrane (Genscript #L00224A60) via the Mini Trans-Blot® Electrophoretic Transfer Cell (Bio0Rad #1703930). Membranes were blocked using 5% milk (w/v) in TBST for 1 hour at room temperature. Primary anti-p-mTOR (Santa Cruz #sc-293133), anti-mTOR (Santa Cruz #sc-517464), anti-Akt (Santa Cruz #sc-5298), anti-PTEN (Cell Signaling, #9559S), and anti-GAPDH (Santa Cruz #sc-47724) primary antibodies, and anti-rabbit Alexa Fluor 647 (Life Technologies #A21245) and anti-mouse Alexa Fluor 647 (Life Technologies #21236) secondary antibodies were used for immunoblotting. Images were captured by Typhoon FLA 9500 (GE Healthcare) and quantification of p-mTOR expression were performed by image J, normalized to the level of GAPDH.

***Quantitative reverse transcription PCR (qRT-PCR)***

CD34^+^ HSPCs pretreated with JNK inhibitor or solution without an inhibitor were co-cultured with MkMPs, non-targeting miR-loaded MkMPs, miR-486-5p inhibitor-loaded MkMPs at 30 MPs per cell, or vehicle control. At 24 hr, cells were harvested and total RNA was isolated, reversed transcribed for qPCR assay as described*^8^*. qPCR assays for *PTEN* and *GAPDH* were performed with PerfeCTa SYBR Green Supermix (QuantaBio) and the following primers: Forward (5’-CGTTACCTGTGTGTGGTGATA -3’), Reverse (5’-CTCTGGTCCTGGTATGAAGAATG-3’) for *PTEN*, and Forward (5’-CCCTTCATTGACCTCAACTACA-3’), Reverse (5’-ATGACAAGCTTCCCGTTCTC-3’) for *GAPDH*. PTEN mRNA level were quantified by normalized to GAPDH mRNA level as the reference gene.

***Intracellular protein analysis by flow cytometry***

100,000 CD34^+^ cells were first pretreated with JNK, Akt, or mTOR inhibitors (or DMSO as control) for 30 min, and co-cultured with MkMPs at 30 MPs/cell, or vehicle control for 16 hours. Cells were fixed in 4% paraformaldehyde for 15 min at room temperature, followed by the permeabilization in 90% methanol for 30 min at 4 ºC. After washing in PBS, cells were stained with Alexa 647-conjugated anti-mTOR (#5048S), PE-conjugated anti-Akt (#8790S), or Alexa 488-conjugated anti-p-Akt (#4071S) antibodies from Cell Signaling, or Alexa 647-conjugated anti-p-mTOR (#564242) from BD bioscience, for 1 hour at room temperature, followed by flow-cytometric analysis.

***Statistical analysis***

Data are presented as means ± standard error of mean (SEM) from at least three replicates. Paired Student’s t test of all data was performed. Statistical significance is defined as **p < 0.05, **p<0.01, ***p<0.001*.

**Supplementary Tables:**

| **Rank** | **MkMPs** | | **PLPs** | |
| --- | --- | --- | --- | --- |
|  | **piRNA ID** | **Fraction** | **piRNA ID** | **Fraction** |
| **1** | hsa_piR_001312 | 19.4 | hsa_piR_001312 | 22.9 |
| **2** | hsa_piR_000765 | 18.4 | hsa_piR_000765 | 21.7 |
| **3** | hsa_piR_020326 | 18.4 | hsa_piR_020326 | 21.7 |
| **4** | hsa_piR_016658 | 10.2 | hsa_piR_017724 | 8.8 |
| **5** | hsa_piR_017724 | 9.1 | hsa_piR_004308 | 3.8 |

**Table S1.** Top 5 expressed human piRNA in MkMPs and PLPs. Piwi-interacting RNAs (piRNAs), which are distinct from miRs, are 24-31 nt in length. As described in Material and Method, we map cDNA sequences (18-40 nt) to human piRNA sequences from the piRNABank (http://pirnabank.ibab.ac.in/). Sequencing data analysis shows that 149 and 152 of a total 458 piRNAs in piRNABank were expressed (average CPM>=1) in MkMPs and PLPs, respectively.. The top 5 expressed piRNAs comprised 76% and 79% of total piRNAs carried by MkMPs and PLPs, respectively.

| **Rank** | **MkMPs** | | **PLPs** | |
| --- | --- | --- | --- | --- |
|  | **snoRNA ID** | **Avg. CPM** | **snoRNA ID** | **Avg. CPM** |
| **1** | SNORD29 | 147576 | SNORD29 | 282862 |
| **2** | SNORD68 | 125014 | SNORD104 | 178695 |
| **3** | SNORD104 | 124403 | SNORD68 | 97625 |
| **4** | SNORD42A | 46614 | SNORD42A | 50157 |
| **5** | SNORD26 | 27796 | SNORD44 | 28201 |
| **6** | SNORD99 | 23774 | SNORD99 | 23767 |
| **7** | SNORD44 | 21651 | SNORD26 | 22674 |
| **8** | SNORD50A | 17159 | SNORD43 | 21394 |
| **9** | SNORD43 | 16061 | SNORD2 | 17408 |
| **10** | SNORD2 | 14935 | SNORD50A | 16937 |

**Table S2.** Top 10 expressed human snoRNA in MkMPs and PLPs. Based on the expression level, among small RNAs of 40-150 nt length, more than 99% of the mapped small RNAs were small nucleolar RNA (snoRNAs)

| **Rank** | **MkMPs** | | | **PLPs** | |
| --- | --- | --- | --- | --- | --- |
|  | **Non-snoRNA ID** | **Avg. CPM** | **Non-snoRNA ID** | | **Avg. CPM** |
| **1** | RNU4ATAC | 1205 | RNU12 | | 933 |
| **2** | RNY5 | 656 | MIR3607 | | 632 |
| **3** | VTRNA1-1 | 600 | RNU4ATAC | | 284 |
| **4** | MIR181B1 | 443 | MIR23A | | 265 |
| **5** | RNU12 | 411 | VTRNA1-1 | | 79 |
| **6** | VTRNA2-1 | 249 | RNY5 | | 51 |
| **7** | MIR3607 | 235 | MIR15A | | 32 |
| **8** | MIR1248 | 147 | SCARNA4 | | 29 |
| **9** | SCARNA4 | 133 | SCARNA11 | | 26 |
| **10** | MIR23A | 80 | RNU11 | | 25 |

**Table S3.** Top 10 expressed human non-snoRNA in MkMPs and PLPs.

| **miR (Gene ID)** | **MkMPs (CPM)** | **Mks (CPM)** | **Fold Change** | **FDR** |
| --- | --- | --- | --- | --- |
| **mir-19b-1//mir-19b-2_3P** | 1999.33 | 378.86 | 2.43 | 0.00025 |
| **mir-181b-1//mir-181b-2** | 1869.99 | 363.50 | 2.37 | 0.00051 |
| **mir-378a_3P** | 1766.13 | 281.71 | 2.92 | 0.00009 |
| **mir-19a_3P** | 986.87 | 170.89 | 2.66 | 0.00012 |
| **mir-30b** | 873.83 | 129.73 | 2.99 | 0.00000 |
| **mir-17** | 452.59 | 101.47 | 2.01 | 0.00172 |
| **mir-335** | 394.39 | 44.40 | 3.93 | 0.00000 |
| **mir-106b** | 388.20 | 70.16 | 2.47 | 0.00001 |
| **mir-345** | 361.60 | 66.55 | 2.49 | 0.00081 |
| **mir-155** | 343.28 | 54.40 | 2.91 | 0.00003 |
| **mir-181c** | 324.84 | 64.18 | 2.29 | 0.00005 |
| **mir-20a** | 288.40 | 60.84 | 2.15 | 0.00059 |
| **mir-320b-1//mir-320b-2_3P** | 266.37 | 47.07 | 2.57 | 0.00027 |
| **mir-451a** | 190.90 | 27.36 | 3.18 | 0.00000 |
| **mir-151b_3P** | 152.06 | 23.24 | 2.89 | 0.00006 |
| **mir-494_3P** | 135.02 | 28.49 | 2.13 | 0.00177 |
| **mir-369_3P** | 121.11 | 23.72 | 2.23 | 0.00539 |
| **mir-376c_3P** | 118.30 | 25.94 | 2.03 | 0.00204 |

**Table S4.** Significantly enriched (*p < 0.01*; differential expression ≥two-fold) miRs (CPM >100) in MkMPs compared to Mks.

**Supplemental Figures**

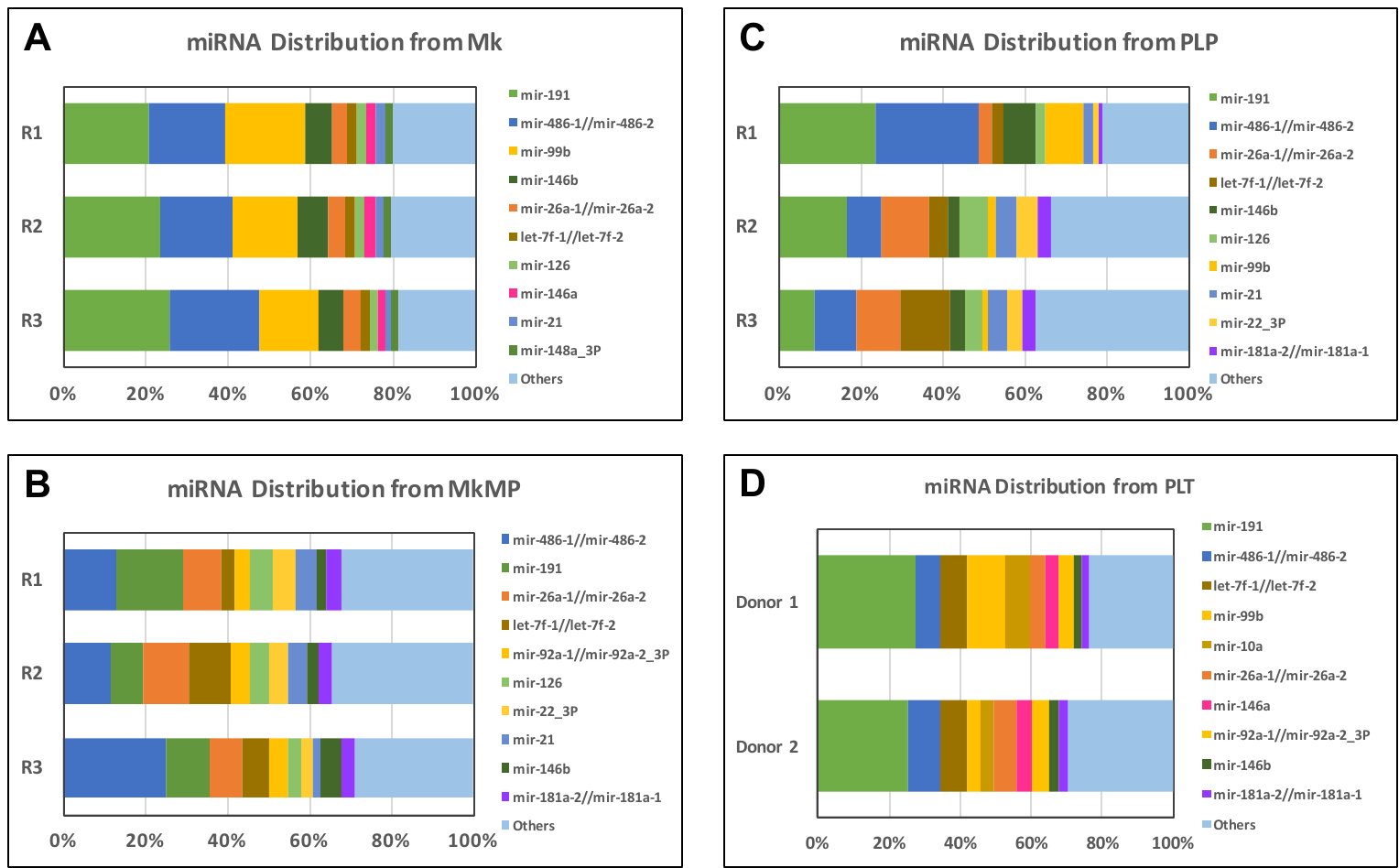

**Figure S1**. Distribution of top 10 miRs from 3-donor (A) Mks, (B) MkMPs, (C) PLPs, and 2 -donor (D) PLTs.

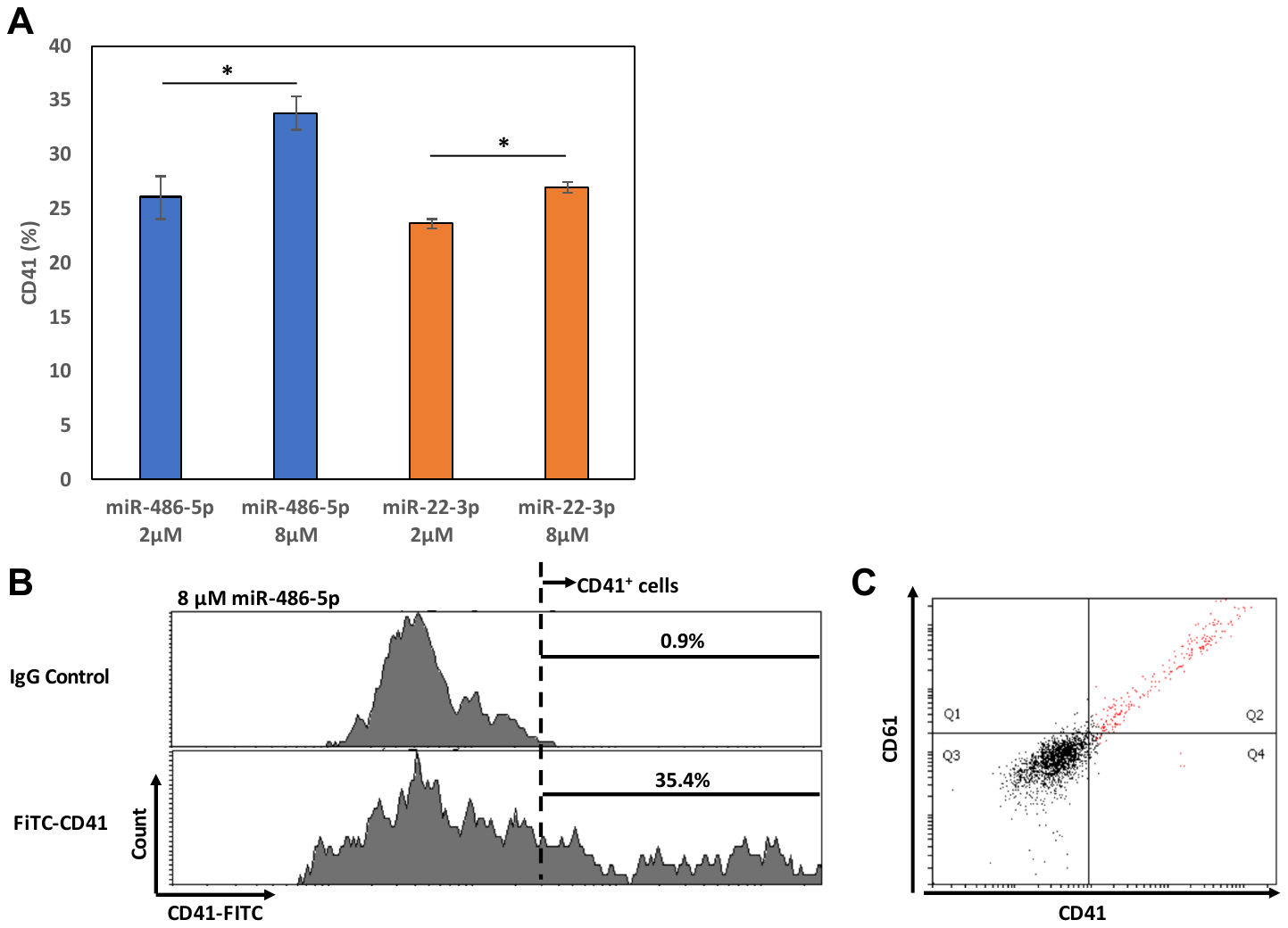

**Figure S2**. 200,000 CD34^+^ HSPCs were transfected with 2 or 8 µM miR-486-5p or miR-22-3p mimics. (A) Cells were harvested for flow cytometric analysis on CD41 expression at day 10. (B) An example of using FITC IgG antibody control to determine the CD41^+^ population in histograms. (C) Cells transfected with 8μM miR-486-5p were also harvested at day 10 for flow cytometric analysis with a quadrant gate for CD41 and CD61 expression. Error bars in (A) represent standard error of mean of 2 biological replicates. **p<0.05*.

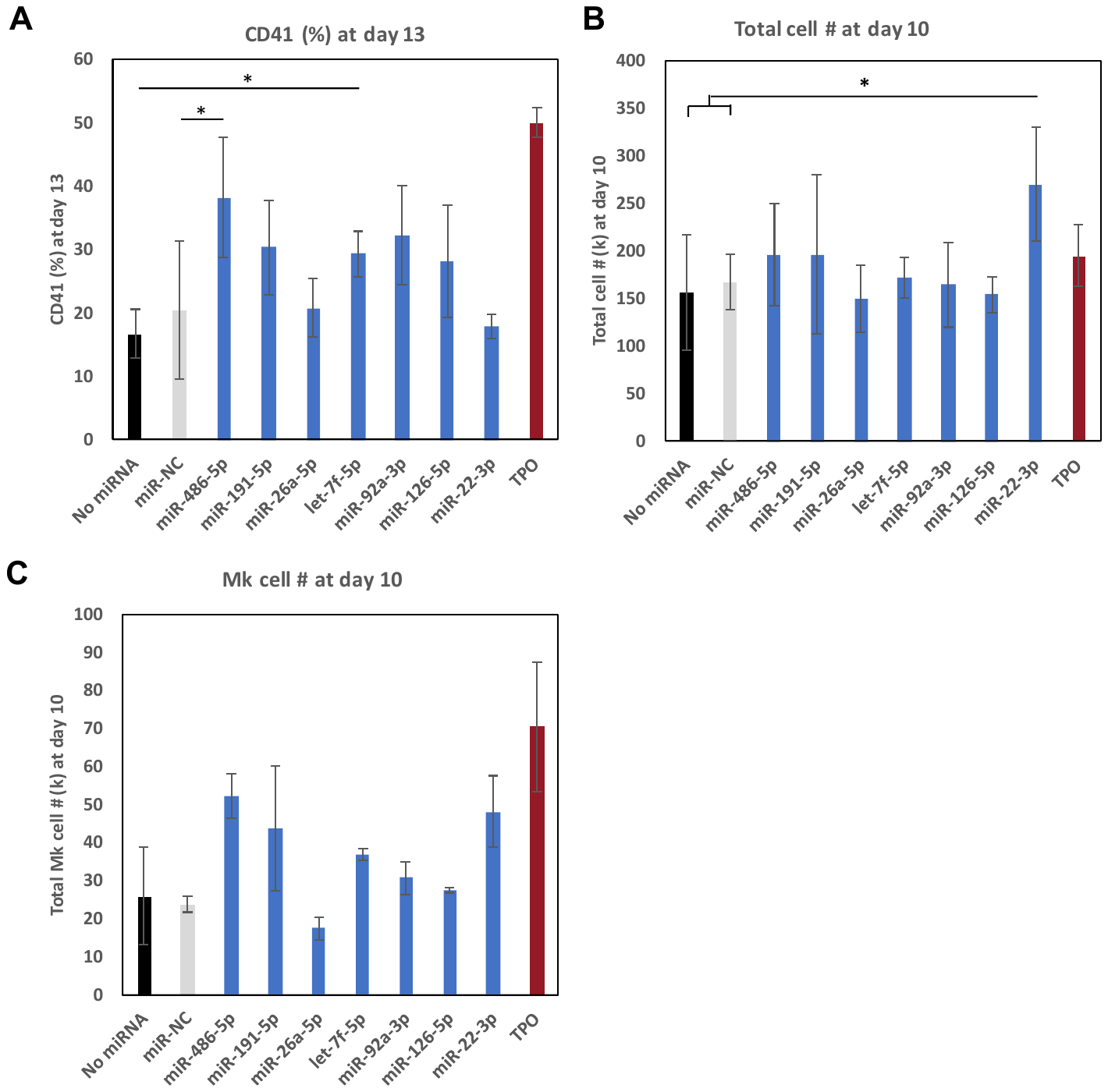

**Figure S3. Effect of single miRs on Mk differentiation.** CD34^+^ HSPCs were transfected with 8 µM miR mimics (N=2), miR negative control (miR-NC), or without miRs (No miR), and cells were cultured in minimal medium (IMDM supplemented with 10% BIT and 50 ng/ml SCF) but without Tpo. Cells cultured in Tpo-supplemented medium (100 ng/ml Tpo) served as positive control (TPO). Cells were harvested for flow cytometric analysis for **(A)** CD41 expression at day 13. **(B)** Total cell counts and **(C)** total Mk-cell counts at day 10. Error bars represent the standard error of mean from 2 biological replicates. **p < 0.05.*

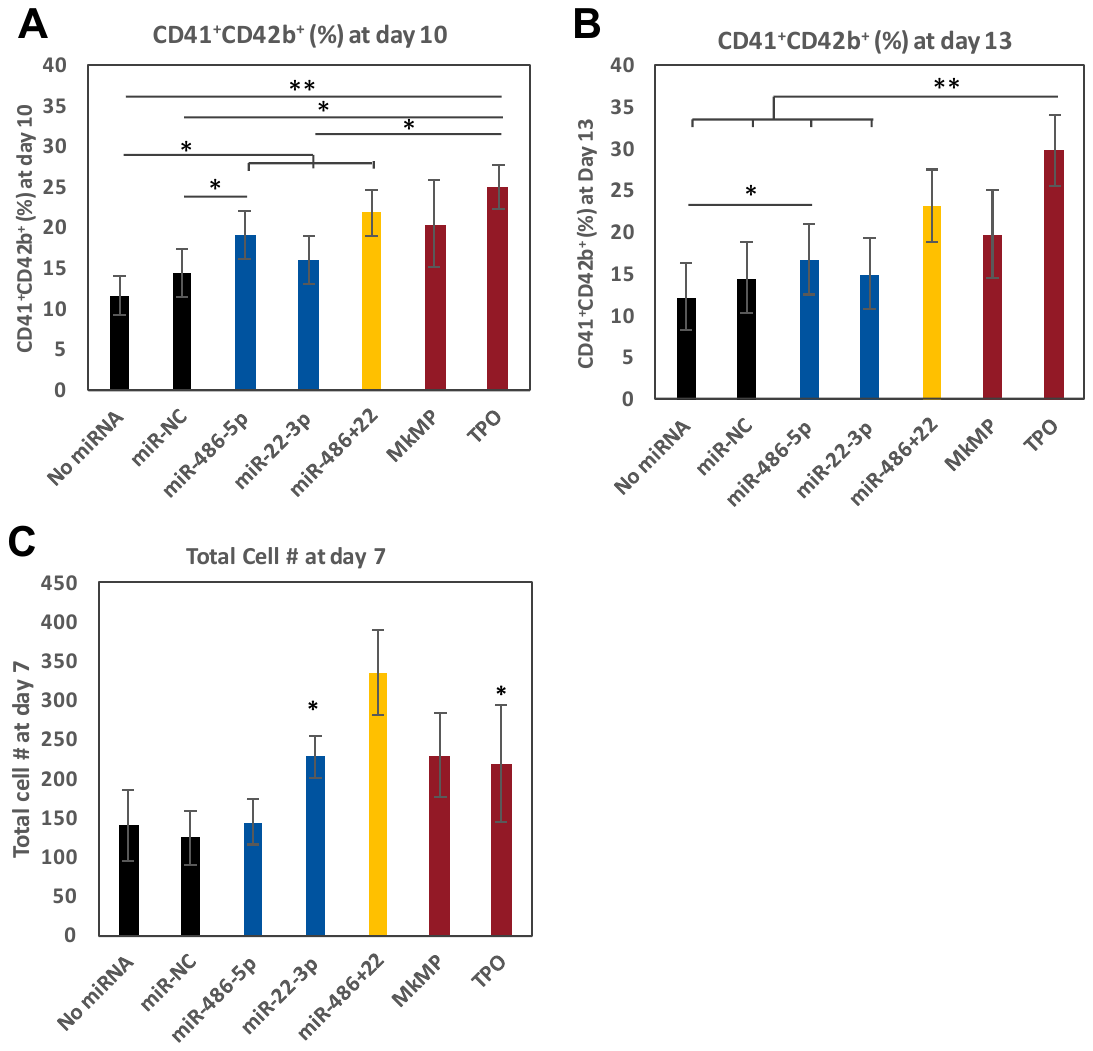

**Figure S4. Effect of single miRs or miR pairs on cell expansion.** CD34^+^ HSPCs were transfected with miR mimics (N=8), miR negative control (miR-NC, N=8), or without miRs (No miR, N=8), and cells were cultured in minimal medium (IMDM supplemented with 10% BIT and 50 ng/ml SCF) without TPO. Cells cultured in TPO-supplemented medium (100 ng/ml TPO, N=6) or cells co-cultured with MkMPs (N=3) served as positive control (TPO, MkMP). The percent of CD41^+^CD42^+^ at day 10 (A) and day 13 (B), and total cell numbers (C) at day 7 were measured by flow cytometry. miR-22-3p significantly promotes cell proliferation. Error bars represent the standard error of mean from 3-8 biological replicates. * represent the comparison to negative controls (No miRNA or miR-NC) unless otherwise indicated on panels A and B. **p < 0.05, **p < 0.01.*

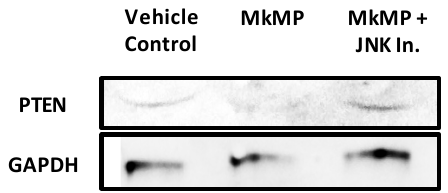

**Figure S5. PTEN expression.** CD34^+^ HSPCs (pretreated with a JNK signaling inhibitor or solution without an inhibitor) were co-cultured with MkMPs. The culture with only CD34^+^ HSPCs served as vehicle control. Cells were harvested after 24 hours of co-culture, and PTEN expression was examined by immunoblotting.
